# Spatiotemporal analysis of axonal autophagosome-lysosome dynamics reveals limited fusion events trigger two-step maturation

**DOI:** 10.1101/2022.02.17.480915

**Authors:** Sydney E. Cason, Saurabh S. Mogre, Erika L.F. Holzbaur, Elena F. Koslover

## Abstract

Macroautophagy is a homeostatic process required to clear cellular waste including aggregated proteins and dysfunctional organelles. Neuronal autophagosomes form constitutively in the distal tip of the axon and are actively transported toward the soma, with cargo degradation initiated en route. Cargo turnover requires autophagosomes to fuse with lysosomes to acquire degradative enzymes; however, the timing and number of these fusion events in the axon have proven difficult to detect using microscopy alone. Here we use a quantitative model, parameterized and validated using data from live and fixed imaging of primary hippocampal neurons, to explore the autophagosome maturation process on a cellular scale. We demonstrate that retrograde autophagosome motility is independent from lysosomal fusion, and that most autophagosomes fuse with only a few lysosomes by the time they reach the soma. Furthermore, our imaging and model results highlight the two-step maturation of the autophagosome: fusion with a lysosome or late endosome is followed by the slow degradation of the autophagosomal inner membrane before actual cargo degradation can occur. Together, rigorous quantitative measurements and mathematical modeling elucidate the dynamics of autophagosome-lysosome interaction and autophagosomal maturation in the axon.

## Introduction

Neurons are terminally differentiated cells that last throughout the lifetime of the organism. One important pathway for maintaining cellular health and homeostasis over this long time period is macroautophagy (hereafter: autophagy), the formation of “self-eating” double-membraned organelles that engulf and degrade cellular waste in order to recycle macromolecular components (Figure 1A) (***Yin et al., 2016***). Defects in neuronal autophagy are implicated in most neurodegenerative disorders including Parkinson’s disease, Alzheimer’s disease, and Amyotrophic Lateral Sclerosis (***Wong and Holzbaur, 2015***). Further, genetically blocking autophagic vacuole (AV) formation causes neurodegeneration in mice (***Hara et al., 2006***; ***Komatsu et al., 2006***). Given the importance of autophagy in the maintenance of neuronal homeostasis, it is essential to gain a quantitative understanding of the pathway, to elucidate how perturbations in rates of autophagy may either negatively or positively affect neuronal health.

**Figure 1.**
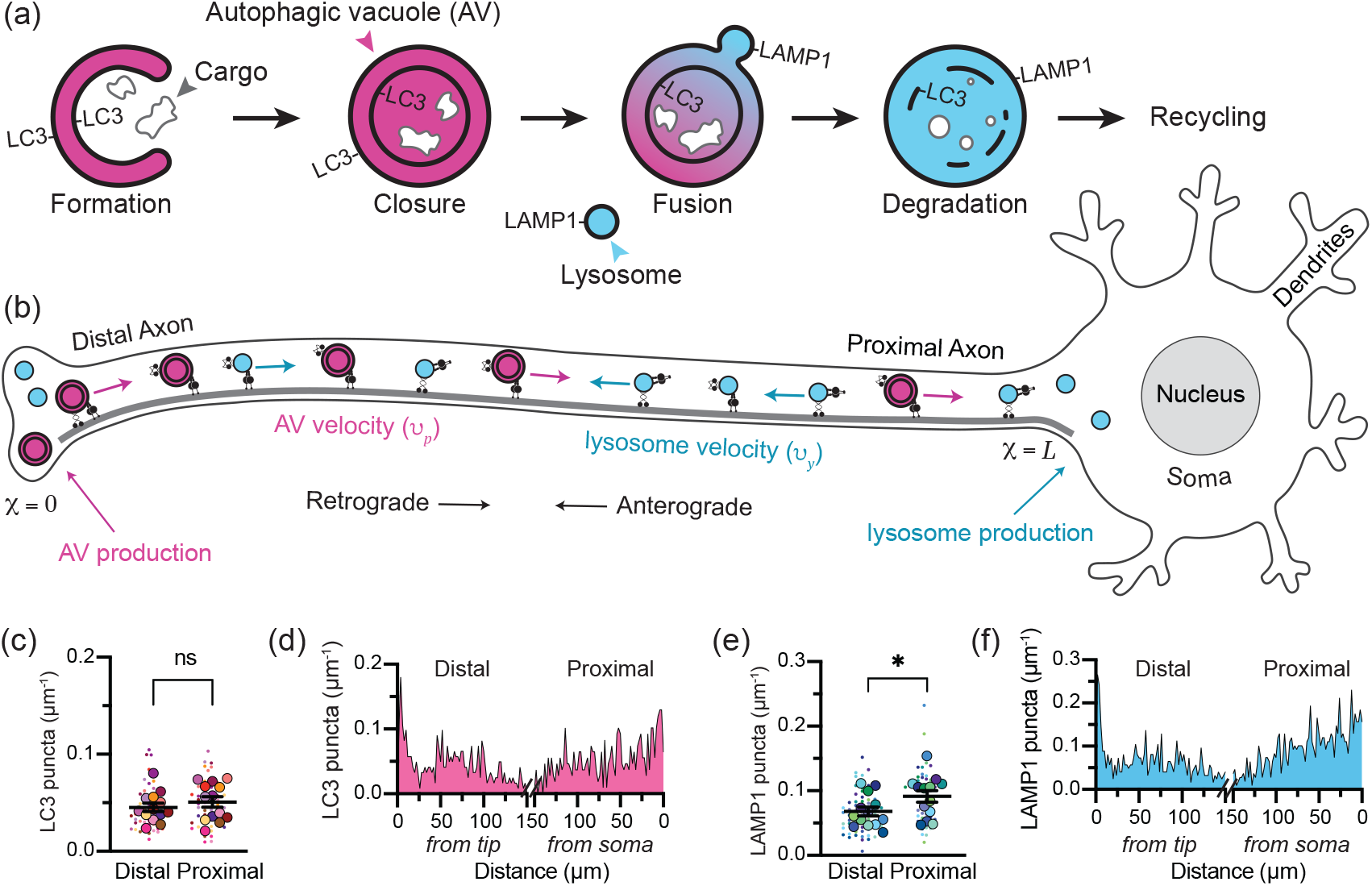
Neuronal autophagosomes form in the distal axon and fuse with lysosomes during transit to the soma. **(a)** Schematic illustrating the autophagy pathway. The developing phagophore engulfs cargo and seals its double membrane to form an autophagic vacuole (AV). The inner and outer membranes are initially decorated with lipidated LC3; however, the LC3 on the outer membrane is cleaved off following closure. AVs fuse with late endosomes and LAMP1-containing lysosomes to acquire degradative enzymes. The autophagic cargo is then broken down and the resulting macromolecules are recycled by the cell. **(b)** Schematic illustrating axonal autophagy. AV biogenesis primarily occurs in the distal tip of the axon, while lysosome biogenesis occurs in the soma. AVs are transported retrograde toward to the soma, while lysosomes move processively in both anterograde and retrograde directions. During this microtubule-based transport, the two organelles encounter one another and have some probability of fusing to facilitate AV maturation. In this study, the distal axon is defined as within 250μm of the axon tip and the proximal axon as within 250μm of the soma, with the total axon length (*x* = *L*) determined experimentally. **(c)** Linear density of endogenous LC3 puncta, detected with RRID:AB_881433 or RRID:AB_11150489. *n* = 13 trials; unpaired t test (*p* = 0.4504). **(d)** Spatial distribution of LC3+ puncta. *n* = 1059 puncta; 2μm bins. **(e)** Linear density of LAMP1 puncta, detected with RRID:AB_1026176 or RRID:AB_2134500. *n* = 13 trials; unpaired t test (*p* = 0.0489). **(f)** Spatial distribution of LAMP1 puncta. *n* = 1720 puncta; 2μm bins. The following figure supplements are available for Figure 1: **Figure 1–Figure supplement 1**. Antibody validation. **Figure 1–Figure supplement 2**. Axon length *in vitro*. **Figure 1–Figure supplement 3**. Axonal AV density in live neurons.

Neuronal AVs form constitutively at synaptic sites and in the distal tip of the axon (Figure 1B),where they clear aged proteins and organelles from the presynaptic region (***Maday et al., 2012***; ***Goldsmith et al., 2022***). However, the vast majority of protein and organelle production occurs in the soma (***Misgeld and Schwarz, 2017***; ***Farfel-Becker et al., 2019***; ***Koltun et al., 2020***). Thus, neuronal AVs must traverse the length of the axon, up to 1m in humans, to recycle their cargo (***Maday and Holzbaur, 2014***; ***Stavoe and Holzbaur, 2019***). AVs acquire molecular motors after formation to drive their transit to the soma (***Fu et al., 2014***; ***Cheng et al., 2015***; ***Cason et al., 2021***). En route, axonal AVs mature by fusing with endolysosomes, organelles containing the digestive enzymes necessary to break down autophagosomal cargo (***Maday et al., 2012***; ***Cason et al., 2021***). Degradatively active endolysosomes, known as lysosomes, are produced in the soma and actively delivered to the axon to fuse with AVs (***Farfel-Becker et al., 2019***; ***Roney et al., 2021***). The maturation of AVs during transport from the axonal tip to the soma is a well-studied phenomenon. However, it has proven experimentally difficult to study the fusion between AVs and endolysosomes along the axon, precluding a quantitative understanding of the spatiotemporal dynamics of maturation.

Mathematical modeling has been used previously to explore the interactions between motile organelles in narrow cellular projections (***Mogre et al., 2020***; ***Agrawal and Koslover, 2021***) such as neuronal axons and fungal hyphae. These studies highlighted the importance of the cross-sectional geometry of the cellular region, as well as organelle production rates and transport dynamics for understanding the interaction probability between organelles (***Agrawal and Koslover, 2021***; ***Williams et al., 2016***; ***Mogre et al., 2020, 2021***). We therefore sought to dissect the mechanisms underlying autophagosomal maturation in the axon by developing a spatially resolved quantitative model of this phenomenon, parameterized from experimental data.

In this work, we construct a comprehensive model of organelle transport, interaction, and maturation during axonal autophagy. The model reproduces features of organelle distribution and maturation observed using endogenous staining and live-cell imaging in primary hippocampal neurons. We incorporate the branched geometry of neuronal axons, and highlight the role of simple parameters including production rates, fusion probability, and motility dynamics. Furthermore, we show that AV maturation is in fact a two-step process, wherein AV-endolysosome fusion is followed by the slow degradation of the inner AV membrane before cargo breakdown can begin. Our quantitative model is used to extract a time-scale for this previously under-appreciated second step of AV maturation. The two-way interplay between experimental measurements and mathematical modeling presented in this work sheds light on the multi-step mechanisms and spatiotemporal distribution of neuronal autophagosome maturation.

## Results

### Autophagic vacuoles mature in the axon under endogenous conditions

Previous studies both *in vitro* and *in vivo* have detected AV maturation by assessing colocalization between fluorescent markers for lysosomes and AVs (***Maday et al., 2012***; ***Stavoe et al., 2016***; ***Hill et al., 2019***; ***Cason et al., 2021***). Most commonly, the protein LC3 (microtubule associated protein 1 light chain 3) or one of its orthologs is used to label AVs. Lysosomes are commonly labeled by lysosome-associated membrane protein 1 (LAMP1); however, LAMP1 also localizes to more immature endocytic compartments and even Golgi-derived carrier vesicles, especially when overexpressed (***Cheng et al., 2018***; ***Farfel-Becker et al., 2019***; ***Lie et al., 2021***). Additionally, we worried exogenous overexpression of LC3 or LAMP1 may affect the quantity of AVs and/or endolysosomes, given their roles in organelle biogenesis and turnover (***Yu et al., 2010***; ***Ma et al., 2012***; ***Shibutani and Yoshimori, 2014***; ***Stavoe et al., 2019***). We therefore performed rigorous immunofluorescence measurements to determine the density and spatial distribution of LC3-containing and LAMP1-containing organelles in the axon under endogenous expression conditions.

In brief, we dissected primary rat hippocampal neurons at embryonic day 18, plated at a low density on glass coverslips, then fixed and permeabilized without detergents after 7-10 days *in vitro*. We then probed for LC3 or LAMP1 using two independent primary antibodies each that were validated both commercially and in house (Figure 1—figure supplement 1). We used a primary antibody against the microtubule-associated protein tau to trace axons (Figure 1—figure supplement 2) and facilitate the identification of proximal (within 250μm of the soma) and distal (within 250μm of the axon tip) axonal segments that did not overlap with adjacent cells. We then acquired Z-stacks at 60*X* magnification and analyzed the resulting maximum projections to quantitate the number of LC3 and LAMP1 puncta per μm of axon.

The number of LC3 puncta in the distal and proximal axon were roughly the same, with an average of 1 punctum every 20μm (≈ 0.05μm^−1^) (Figure 1C). The linear density detected using immunofluorescence is consistent with what we detect using fluorescently-tagged LC3 in the axons of live primary hippocampal neurons and live iPSC-derived neurons (Figure 1—figure supplement 3), as well as published studies quantifying LC3 in the axons of live primary dorsal root ganglia (DRG) neurons (≈ 0.05μm^−1^) (***Maday et al., 2012***) and primary cortical neurons (≈ 0.06μm^−1^) (***Lee et al., 2011***). The distribution of LC3 in the distal axon showed a mildly higher density in the tip/growth cone and a similar density along the proximal axon (Figure 1D).

LAMP1 puncta were slightly more dense, with the number of LAMP1 puncta marginally higher in the proximal axon (1 punctum every ≈ 10μm; ≈ 0.09μm^−1^) than the distal axon (1 punctum every ≈ 15μm; ≈ 0.07μm^−1^) (Figure 1E). The LAMP1 puncta density detected using immunofluorescence is consistent with published studies quantifying LAMP1 puncta in live hippocampal (≈ 0.08μm^−1^) (***Boecker et al., 2020***) or iPSC-derived i^3^ neurons (≈ 0.08μm^−1^) (***Boecker et al., 2020***), and slightly lower than that seen using fluorogenic lysosomal enzyme activity sensors in DRG neurons (≈ 0.14μm^−1^) (***Farfel-Becker et al., 2019***). In the distal region, we observed some accumulation of LAMP1 puncta in the axon tip/growth cone (Figure 1F).

### Endogenous LAMP1 localizes to degradative compartments

Some recent studies have proposed that only a small fraction of axonal lysosomes are degradatively competent (***Gowrishankar et al., 2015***; ***Cheng et al., 2018***; ***Farfel-Becker et al., 2019***). Studies in non-neuronal cells have also proposed a lysosomal activity gradient wherein lysosomes closer to the nucleus are more mature and proteolytically active than those farther from the nucleus (***Johnson et al., 2016***; ***Ferguson, 2018***). We therefore examined the degradative capacity of lysosomes along the axon by measuring the colocalization between endogenous LAMP1 and the endogenous lysosomal enzymes asparagine endopeptidase (AEP) and Cathepsin L (CTSL) (Figure 2A-D). Across the axon, roughly three-quarters of the LAMP1 colocalized with lysosomal enzymes. Lysosomal proteases require low pH to function, so we probed for the presence of the lysosomal vATPase, which pumps protons across the membrane to achieve and maintain the lysosome’s characteristic low pH (≈4.8) (***Johnson et al., 2016***). The vATPase subunit V1 is cytoplasmic and forms an activated vATPase when interacting with the transmembrane V0 subunit. Therefore colocalization between the V1 subunit (ATP6V1F) and membrane-bound LC3 suggests the formation of an active vATPase on the AV membrane. Again, we saw about 80% colocalization in both the proximal and distal axon (Figure 2E,F). These data suggest that the population of LAMP1-positive (LAMP1+) lysosomes along the axon is primarily mature and degradatively competent.

**Figure 2.**
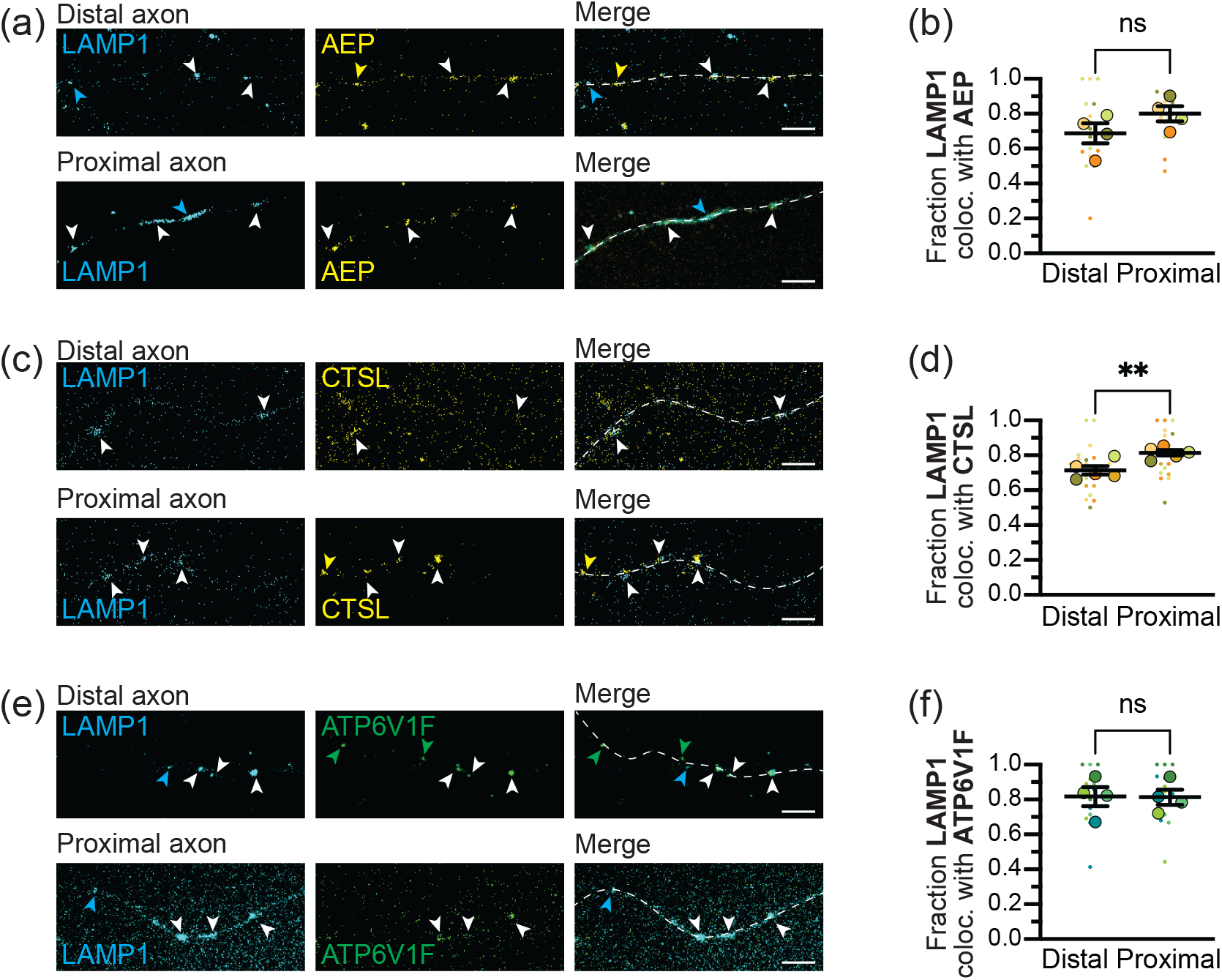
Lysosomes throughout the axon are degradatively competent. Maximum projections **(a)** and quantification **(b)** showing LAMP1 and asparagine endopeptidase (AEP) overlap in the distal and proximal axon. *n* = 4 trials; unpaired t test (*p* = 0.1684). Maximum projections **(c)** and quantification **(d)** showing LAMP1 and Cathepsin L (CTSL) overlap in the distal and proximal axon. *n* = 5 trials; unpaired t test (*p* = 0.0073). Maximum projections **(e)** and quantification **(f)** showing LAMP1 and V-type proton ATPase subunit F (ATP6V1F) overlap in the distal and proximal axon. *n* = 4 trials; unpaired t test (*p* = 0.9588). Dashed line represents axon. Cyan arrows, LAMP1 alone; yellow/green arrows, respective marker alone; white arrows, colocalization. Scale bar, 5 μm. Fractions are all over the total LAMP1+ puncta in that region. Bars throughout show mean ± SEM. Dashed line represents axon. ns, p > 0.05; **, p < 0.01.

### Most AVs mature along the axon

Next, we quantified colocalization between LC3 and LAMP1 and found that roughly 50% of the AVs in the distal axon colocalized with LAMP1 (Figure 3A,B). Considering our optical resolution (200nm) these colocalized puncta likely represent fused or fusing organelles. A higher number of AVs were positive for LAMP1 in the proximal axon; thus, an additional ≈25% of AVs fused with a LAMP1+ organelle in the axon shaft during transit to the soma (Figure 3A,B). About 30% of the LAMP1 across the axon colocalized with LC3, and there was no difference between the distal and proximal axon (Figure 3–figure supplement 1).

**Figure 3.**
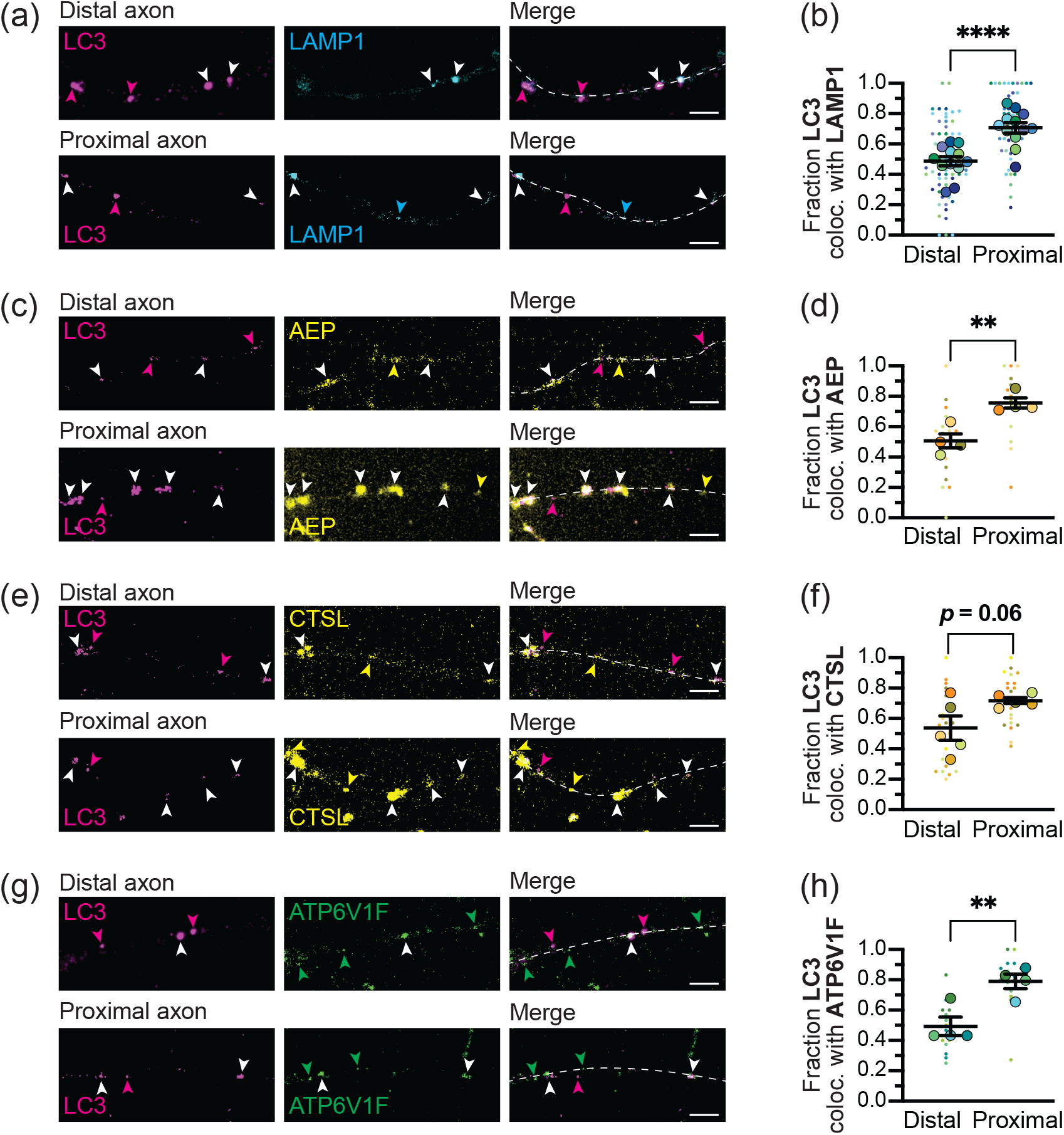
Spatial distribution of LC3 colocalization with lysosomal markers under endogenous conditions. Maximum projections **(a)** and quantification **(b)** showing LC3 and LAMP1 overlap in the distal and proximal axon. *n* = 12 trials; unpaired t test (*p* < 0.0001). Maximum projections **(c)** and quantification **(d)** showing LC3 and AEP overlap in the distal and proximal axon. *n* = 4 trials; unpaired t test (*p* = 0.0045). Maximum projections **(e)** and quantification **(f)** showing LC3 and CTSL overlap in the distal and proximal axon. *n* = 5 trials; unpaired t test (*p* = 0.0605). Maximum projections **(g)** and quantification **(h)** showing LC3 and ATP6V1F overlap in the distal and proximal axon. *n* = 4 trials; unpaired t test (*p* = 0.0090). Magenta arrows, LC3 alone; cyan/yellow/green arrows, respective lysosomal marker alone; white arrows, colocalization. Scale bar, 5 μm. Fractions are all over the total LC3 puncta in that region. Bars throughout show mean ± SEM. Dashed line represents axon. ns, p > 0.05; *, p < 0.05; **, p < 0.01; ****, p < 0.0001. The following figure supplement is available for Figure 2: **Figure 3–Figure supplement 1**. LAMP1 puncta colocalized with LC3.

To determine whether LC3-positive (LC3+) organelles were fusing with degradative lysosomes, we assessed colocalization with endogenous lysosomal enzymes AEP and CTSL. Roughly half of the LC3 puncta in the distal axon colocalized with lysosomal enzymes (Figure 3C-F). Colocalization with either enzyme increased in the proximal axon, indicating further fusion of AVs with endolysosomes during translocation toward the soma. We also looked for colocalization of endogenous LC3 with the V1 subunit of the vATPase, a marker of active proton pumps. Again, roughly half of the LC3 puncta in the distal axon colocalized with ATP6V1F, and an additional ≈30% of AVs appeared to acquire ATP6V1F during transit along the axon shaft (Figure 3G,H). We therefore conclude that half of the axonal AV population fuses with a degradatively active lysosome prior to leaving the distal axon, while an additional quarter of the axonal AV population fuses with an active lysosome along the mid-axon prior to reaching the proximal axon and soma.

### Mathematical modeling elucidates interplay of transport and fusion in autophagosomelysosome distributions

We next proceeded to develop a coarse-grained mathematical model for axonal AV maturation through fusion with endolysosomes. The model is parameterized against experimental data and aims to elucidate how organelle transport and interaction parameters dictate the spatial distribution of lysosomes, AVs, and fusion events. Because axons are much longer than they are wide, we simplify the model system to a one-dimensional domain of length *L* = 1055μm, representing the average length of primary hippocampal axons at 7-10 days *in vitro* (Figure 1—figure supplement 2). The model includes the biogenesis of AVs in the distal axon tip (***Maday and Holzbaur, 2014***) and the production of lysosomes in the soma (***Farfel-Becker et al., 2019***), along with switches between different motility states (***Fu et al., 2014***; ***Cason et al., 2021***) and AV-lysosome fusion events (Figure 4). We explore organelle distributions both with stochastic agent-based simulations of discrete particles and in a mean-field sense, by solving for the continuous spatial densities of different organelle states.

**Figure 4.**
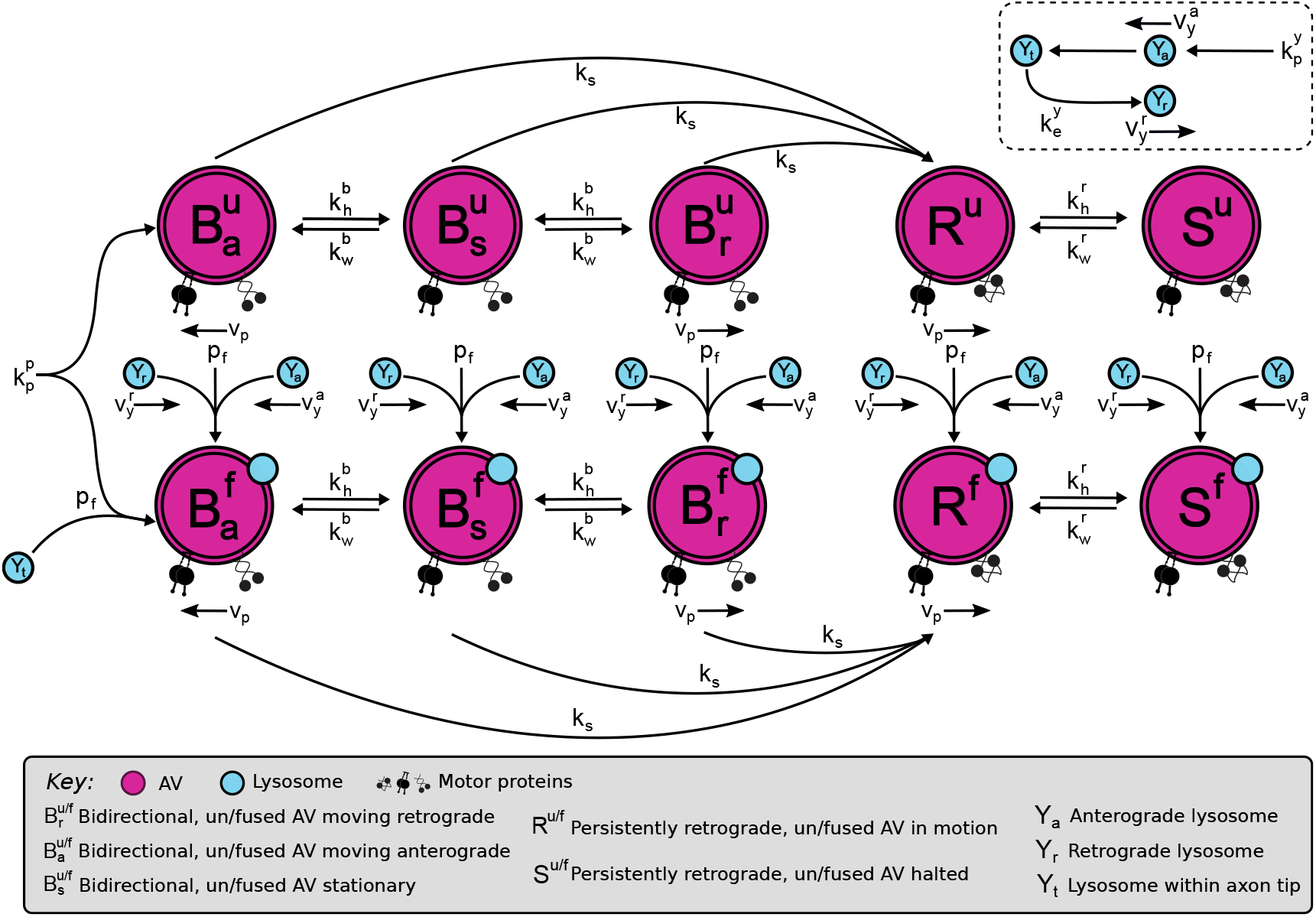
Mathematical model describes AV motility and endolysosome fusion as interconverting states. Magenta states indicate AVs, with top row corresponding to AVs that have not fused with an endolysosome and are bidirectional (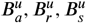 depending on current motion anterograde, retrograde, or stationary, respectively) or persistently retrograde (*R*^*u*^: in motion, *S*^*u*^: paused). Bottom row shows corresponding states for AVs that have fused with an endolysosome. Lysosome states (*Y*_*a*_: anterograde, *Y*_*r*_: retrograde, *Y*_*t*_: tip localized) are shown in cyan. Transitions between states are marked by arrows with the corresponding transition rates labeled.

#### Model for AV transport and distribution

In the model, AVs are formed at the distal axon tip (*x* = 0) at rate 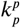. Nascent AVs engage in short bidirectional movements, or remain relatively stationary (***Fu et al., 2014***; ***Cason et al., 2021***). In the distal axon, the majority (roughly 80%) of LC3+ AVs move less than 10μm over the course of a 1-3 minute video acquisition (Figure 5A). Such puncta are either completely stationary or in a bidirectional state consisting of short anterograde or retrograde runs interspersed with stationary periods (Figure 5B). The pause-free velocity in the bidirectional state is set to *v*_*p*_ = 0.75*μm/s* in both directions (***Boecker et al., 2020***). The spatial densities of retrograde, anterograde, and stationary AVs in the bidirectional state are described by *B*_*r*_(*x*), *B*_*a*_(*x*), *B*_*s*_(*x*).

**Figure 5.**
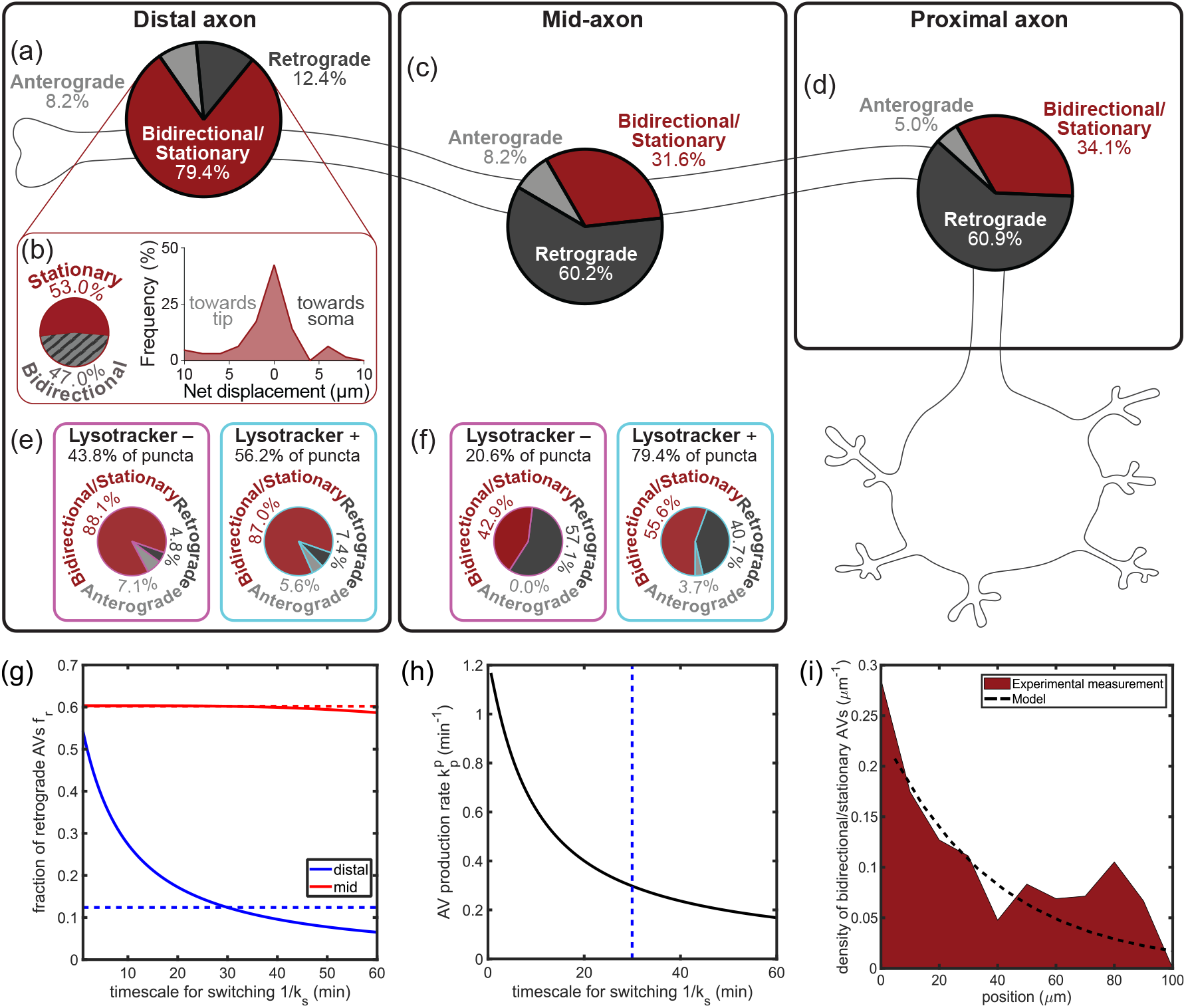
Production and motility switching rates determine spatial densities of bidirectional and retrograde AVs. **(a)** Motility of mCherry-LC3+ puncta in the distal axon (*n* = 36 cells). The majority of puncta exhibited nonprocessive motility (< 10μm net displacement over the course of a 1-3 minute video), with < 15% moving anterograde (≥ 10μm towards the tip) or retrograde (≥ 10μm towards the soma). **(b)** Among the non-processive puncta (from *n* = 13 cells), approximately half are classified as stationary, with an overall trajectory range < 3*μ*m. The remaining puncta are classified as bidirectional. **(c)** The displacement distribution among non-processive puncta is approximately symmetric, indicating unbiased motion. **(d-e)** Motility states of mCherry-LC3+ puncta within the mid- (**b**, *n* = 40 cells) or proximal (**c**, *n* = 35 cells) axon. The majority of puncta exhibited retrograde (≥ 10μm towards the soma) motility. **(f-g)** Motility states of mCherry-LC3+ puncta within the distal **(d)** or mid-axon **(e)**, separated based on the fusion state determined by colocalization with LysoTracker. The retrograde moving fraction (≥ 10μm net displacement) among fused and unfused AVs was not significantly different within the distal (*n* = 14 cells, *p*=0.6933, Fisher’s exact test) or mid-axon (*n* = 7 cells, *p* = 0.6722, Fisher’s exact test). **(h)** From quantitative modeling, predicted fraction of AVs exhibiting retrograde motility within the distal (blue) and the mid axon (red), plotted against the timescale for switching (*τ*_*s*_ = 1/*k*_*s*_). The observed fractions within hippocampal axons are denoted by the corresponding dashed lines. **(i)** Model AV production rate 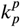 required to achieve the measured LC3+ density in the distal axon, plotted against the timescale for switching. The dashed line denotes the switching time obtained in **(h)** by fitting the retrograde fraction in the distal region. **(i)** Distribution of stationary/bidirectional AVs (with ≤ 10*μ*m net displacement) in the distal axon. The dashed black line denotes the distribution predicted by the mathematical model (*B*_*s*_ + *B*_*r*_ + *B*_*a*_ + *S*).

We subdivide this population of stationary/bidirectional AVs (with < 10μm displacements) further by classifying as stationary all particles that have a range < 3μm during the 1-3 minute imaging period. This gives the fraction stationary among the non-processive population as *f*_*s*_ ≈ 53% (Figure 5B). The remaining particles are classified as bidirectional. The average run-length in a consistent direction for such particles is estimated as *λ* = 2*μ*m, corresponding to a rate constant 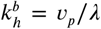 for transition from either the anterograde or retrograde into a stationary state. Given the approximately symmetric histogram of the AV displacements for the bidirectional/stationary population (Figure 5B), we assume that the bidirectional motion is largely unbiased, with rate constant 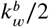 for transitioning from a stationary state into either an anterograde or retrograde run. The restarting rate is estimated according to the observed stationary fraction, as 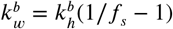.

All bidirectional AVs have a constant rate *k*_*s*_ of switching to a processive retrograde motile state [density *R*(*x*)] with constant velocity *v*_*p*_. While only 12.4% of LC3+ puncta are retrograde in the distal axon (Figure 5A), this fraction rises to ≈ 60% in the mid-axon and proximal regions (Figure 5C,D), with concomitantly fewer bidirectional and stationary particles observed in those regions. Past studies of axonal autophagosome dynamics suggested that fusion with an endolysosome was a prerequisite for switching to retrograde motility (***Cheng et al., 2015***). We analyzed the motility of AVs which did (LysoTracker+) or did not (LysoTracker–) colocalize with LysoTracker, a dye that labels acidified compartments and can therefore be used as a proxy for fusion with endolysosomes. In both the distal and mid-axon, LysoTracker+ AVs were no more likely to exhibit retrograde motion than LysoTracker– AVs, implying the motility switch is not connected to fusion (Figure 5E,F). To minimize model parameters, we therefore assume a single constant switching rate regardless of whether an AV has fused with an endolysosome.

Given our model assumes AVs are produced at the distal tip, we would expect that most LC3+ puncta found in the mid- and proximal axon must have arrived there after undergoing the switch to a retrograde moving state. To account for the remaining stationary puncta observed in these regions, we assume that a processively retrograde AV can switch into a temporary paused state with rate 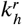 and can resume its retrograde motion with rate 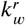. The density of such paused Avs is defined by *S*(*x*). The model is insensitive to the absolute rates 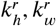. However, the ratio 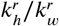 sets the ratio for paused to retrograde AVs (*S*(*x*)/*R*(*x*)) throughout the mid- and proximal axon.

The mean-field model comprises a set of steady-state equations for the densities of AVs in each state:

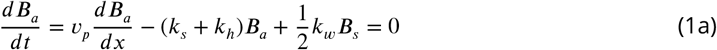

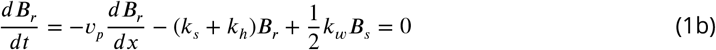

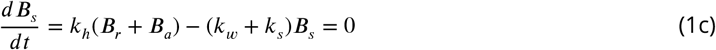

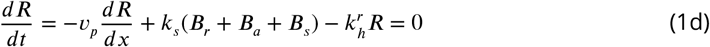

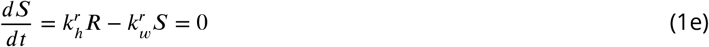

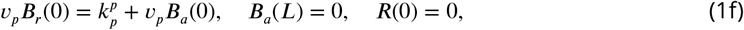

where the last trio of equations describes boundary conditions at the distal tip and the cell body. These linear, homogeneous, first-order equations can be solved using standard matrix methods. We note that the AV production rate 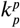 scales the total density of AVs in the domain and does not affect the relative fraction in each state. We can thus directly fit the remaining unknown parameters 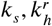 by considering the fraction of AVs in the processively retrograde state in the distal and mid-axonal regions, respectively. Specifically, we compute the fraction of retrograde AVs among those in the distal region (*x* < 250μm) of the linear domain, as a function of the switching rate *k*_*s*_ (Figure 5F). We compare this value to the experimentally measured fraction retrograde among distal AVs (≈ 0.124 ± 0.024, pooled across *n* = 36 cells), defined as puncta that move in the retrograde direction for > 10μm. The comparison allows us to extract a fitted parameter *k*_*s*_ ≈ 0.04min^−1^. In the absence of AV pausing after they enter the retrograde state, we would expect the fraction retrograde in the mid-axonal region to be close to 100%. We adjust the pausing rate to match the experimentally observed mid-axon fraction retrograde of 60.2% (Figure 5C), setting the ratio 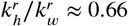.

The AV production rate can be fitted by scaling the average total density of AVs in the distal region to match the experimentally measured value *ρ*_*p*_ = 0.045 ± 0.004μm^−1^ (Figure 1C). The fitted value (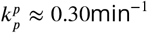; Figure 5H) is within the range previously reported in different neuronal cell types (0.12 − 0.6 min^−1^) (***Maday and Holzbaur, 2014***)].

The mathematical model predicts that the density of bidirectional autophagosomes should fall off with distance away from the distal tip. The length-scale for this decrease depends on the rate *k*_*s*_ of switching into the processive retrograde state, as well as the stopping and restarting rates for bidirectional AVs. In Figure 5I we show that the model predictions are approximately consistent with the observed distal distribution of stationary and bidirectional AVs.

#### Model for autophagosome-lysosome fusion

To explore fusion behavior, the distribution of lysosomes needs to be incorporated into the model. We assume lysosomes are produced in the soma (*x* = *L*) with rate 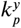. Upon biogenesis, lysosomes enter the axon and move in the anterograde direction towards the axonal tip, at an effective velocity of 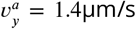. This effective velocity encompasses both the measured pause-free speed of anterograde lysosomes in hippocampal neurons and the frequency of pauses (***Boecker et al., 2020***). Lysosomes that reach the distal tip of the domain (*x* = 0) without fusing with an AV enter a halted state. From there, they have a constant rate 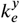 of initiating retrograde motion towards the soma at an effective velocity of 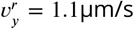, thus exiting the distal tip (Figure. 6A). The densities of anterograde and retrograde lysosomes are defined as *Y*_*a*_(*x*) and *Y*_*r*_(*x*), respectively. The halted state encompasses all lysosomes accumulated in the distal bud of the axon (*Y*_*t*_) without resolving the precise spatial position within that distal bud.

**Figure 6.**
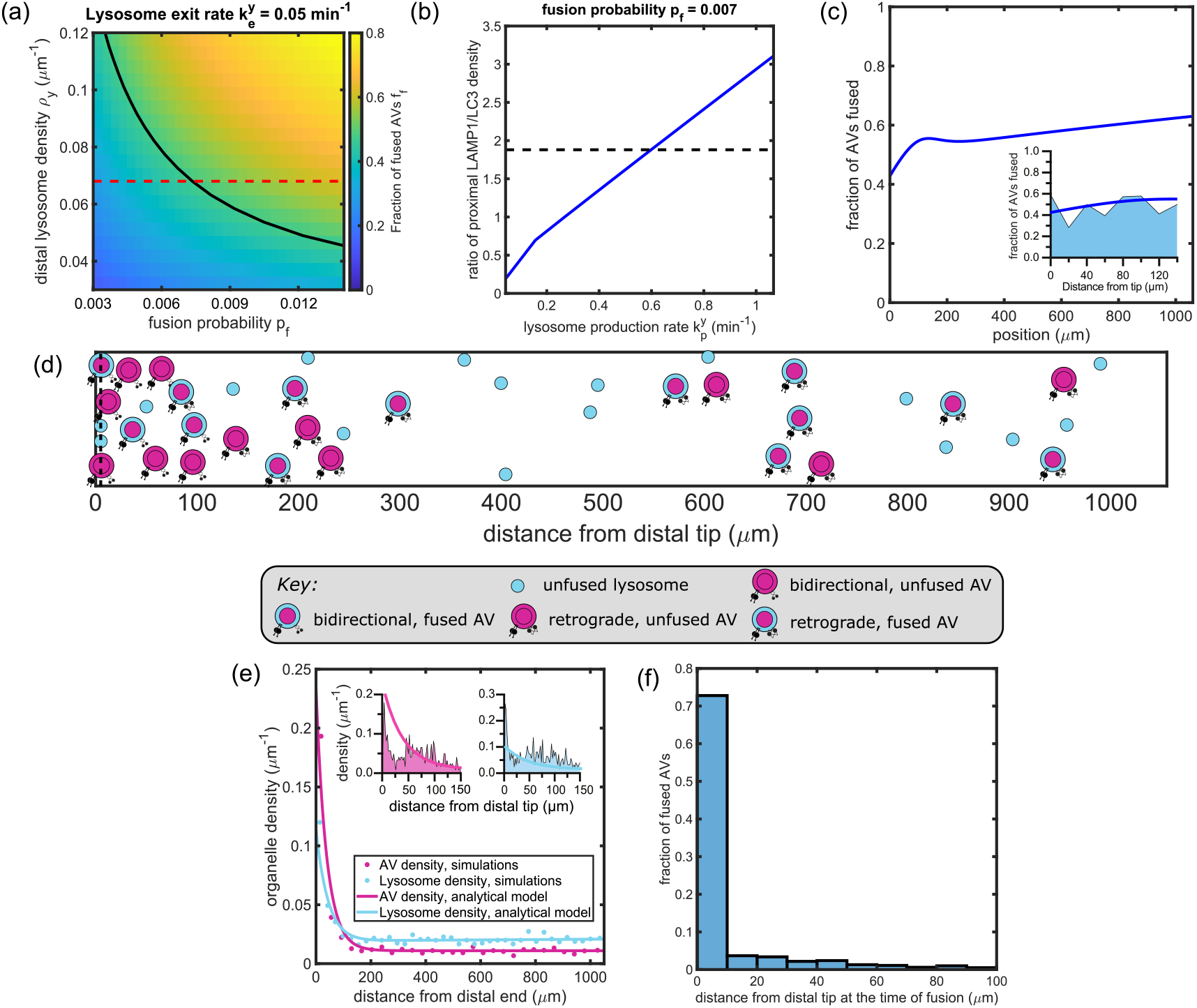
Model for AV-endolysosome interaction dynamics predicts spatial distributions of fused and unfused organelles. **(a)** Fraction of AVs fused within the distal axon *f*_*f*_, plotted against the fusion probability *p*_*f*_, and the lysosome density in the distal region *ρ*_*y*_. The tip-exit rate for lysosomes 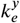 is set to 0.05 per minute. The solid black line denotes observed value of *f*_*f*_ based on LC3+ puncta colocalized with LAMP1+ puncta in the distal axon. The dashed red line denotes the observed density of LAMP1+ puncta in the distal axon. **(b)** The ratio of the linear density of lysosomes to AVs in the proximal axon, plotted against the lysosome production rate. The dashed black line denotes the measured value determined by enumerating LAMP1+ and LC3+ puncta in the proximal axon. **(c)** Spatial variation in the fraction of AVs fused at different positions along the axon. The inset zooms into the distal region, overlaid with the experimentally observed distribution obtained by enumerating LC3+ puncta colocalized with LAMP1. **(d)** Snapshot from agent-based simulation of organelle dynamics, after reaching steady-state. For clarity, organelle size and axon cross-section (vertical axis) is not shown to scale. Video of simulation is provided as Figure 6 – Supplemental Video 1. **(e)** The linear density of LC3+ puncta (magenta) and LAMP1+ puncta (cyan) along the axon. Solid lines are obtained from mean-field model, and dots from stochastic simulations. Insets show comparison to experimentally measured densities in distal region, from Fig. 1(d,f). **(f)** Histogram of position at first fusion for individual AVs, extracted from simulated trajectories. Vertical axis is normalized to the overall number of AVs that undergo fusion before reaching the soma. The most distal 100*μ*m are shown.

We assume a constant probability of fusion *p*_*f*_ each time an AV and a lysosomal particle pass each other. Upon fusion, the endolysosome disappears, and the autophagosome is marked as fused. In our initial model, we assume that the ability of the AV to fuse with subsequent endolysosomes is lost after the initial fusion event. This is consistent with a model wherein the fusion machinery is inhibited following fusion (***Saleeb et al., 2019***).

For a given retrograde-moving AV at position *x*, the flux of anterograde-moving lysosomes passing by it per unit time can be expressed as 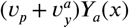, where the prefactor is the relative velocity of the two particles. Similarly, the flux of retrograde lysosomes passing by a retrograde AV is given by 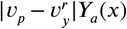, and analogous expressions can be formed for all combinations of lysosome motility (anterograde or retrograde) and AV motility (anterograde, retrograde, or stationary). The rate at which fusion occurs is proportional to this flux multiplied by the fusion probability. We therefore write the mean-field equations for unfused AV densities:

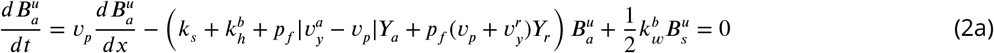

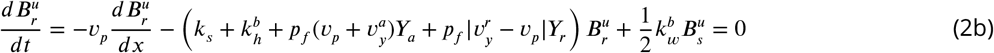

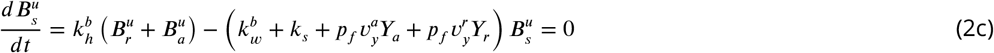

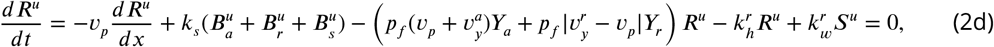

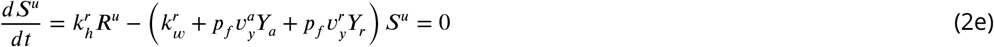

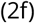

where 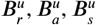 are densities of unfused AVs in the bidirectional retrograde, anterograde, and stationary states, respectively. *R*^*u*^ and *S*^*u*^ denote densities of unfused AVs in the persistently retrograde state that are currently moving or paused, respectively. The corresponding equations for lysosome densities are given by

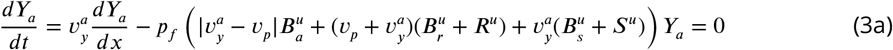

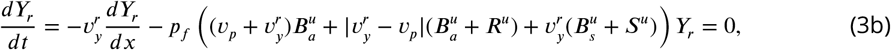

where *Y*_*a*_ and *Y*_*r*_ are the densities of lysosomes moving in the anterograde and retrograde direction, respectively. For this system of equations, the boundary conditions are:

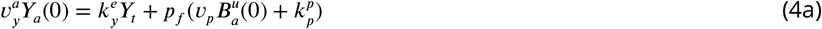

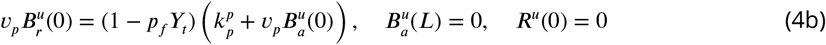

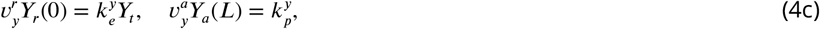

where the first equation gives the steady-state condition for lysosome halted at the distal tip.

This set of non-linear equations is solved numerically as described in the Methods to obtain the distributions of lysosomes and unfused AVs. The total density of fused AVs, corresponding to puncta labeled with both LC3 and LAMP1, can be found as 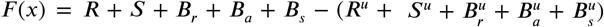. The total density of LAMP1+ puncta is *Y* (*x*) = *Y*_*a*_ + *Y*_*r*_ + *F*. We fit three additional model parameters pertaining to fusion and lysosome behavior 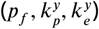 by matching three different metrics to experimentally observed data: the average density of LAMP1+ puncta in the distal region (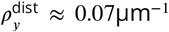; Figure 1E), the fraction of AVs that have fused with a lysosome in the distal region 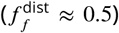 as measured by colocalization with LAMP1 (Figure 3B), and the ratio of LAMP1+ puncta to LC3+ puncta in the proximal region (*R*^prox^ ≈ 1.9; Figure 1C,E). The distal and proximal regions are defined as segments of length 250μm from the distal and proximal ends of the domain, respectively. The distal lysosomal density also includes those lysosomes halted in the distal tip. The resulting fitted parameter values are listed in Table 1.

**Table 1.** Parameters for modeling organelle dynamics in autophagy

| Fixed parameters |  |  |  |  |  |
| --- | --- | --- | --- | --- | --- |
| Parameter | Description | Value | Reference |  |  |
| $L$ | Length of the main axon | 1055$\mu\text{m}$ | Figure 1 S2 | | |
| $v_p$ | AV velocity | 0.75$\mu\text{m/s}$ | *Boecker et al. (2020)* | | |
| $v_y^a$ | Lysosome velocity (anterograde) | 1.3$\mu\text{m/s}$ | *Boecker et al. (2020)* | | |
| $v_y^r$ | Lysosome velocity (retrograde) | 1.1$\mu\text{m/s}$ | *Boecker et al. (2020)* | | |
| Measurements |  |  |  |  |  |
| Parameter | Description | Value | Reference |  |  |
| $\rho_p$ | Distal LC3 density | (0.045±0.004) $\mu\text{m}^{-1}$ | Figure 1c | | |
| $\rho_y$ | Distal LAMP1 density | (0.068±0.007) $\mu\text{m}^{-1}$ | Figure 1e | | |
| $R$ | Proximal LAMP1/LC3 ratio | 1.88±0.11 | Figure 1c,e | | |
| $f_f$ | Distal fused AV fraction | 0.49±0.03 | Figure 3b | | |
| $f_r$ | Distal retrograde AV fraction | 0.12±0.02 | Figure 5a | | |
| Fitted parameters |  |  |  |  |  |
| Parameter | Description | Value (linear) | Reference | Value (branched) | Reference |
| $k_s$ | AV Retrograde switch rate | 0.03min$^{-1}$ | Figure 5g | 0.03min$^{-1}$ | Figure 7b |
| $k_h^r/k_w^r$ | Retrograde AV halt rate (scaled) | 0.66 | – | 0.60 | – |
| $k_p^p$ | AV production rate | 0.30min$^{-1}$ | Figure 5h | 0.30min$^{-1}$ | Figure 7c |
| $p_f$ | Fusion probability | 0.007 | Figure 6a | 0.007 | Figure 7d |
| $k_p^y$ | Lysosome production rate | 0.60min$^{-1}$ | Figure 6b | 3.37min$^{-1}$ | Figure 7e |
| $k_e^y$ | Lysosome tip exit rate | 0.05min$^{-1}$ | Figure 6a | 0.04min$^{-1}$ | Figure 7d |

The distal fraction fused 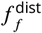 is largely dependent on the fusion probability and the overall density of lysosomes in the distal region, regardless of the combination of production and exit rate that yields that density. We plot this fraction fused in Figure 6A, showing that a unique, low value of the fusion probability *p*_*f*_ ≈ 0.007 matches both observations of the distal lysosomal density and the fraction of fused AVs in the distal region. The distal density of lysosomes is determined by the balance between the lysosomal production rate and their retrograde exit rate from the distal tip. A fixed value of this distal density can be achieved by low production but long pausing at the tip (low 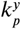 and 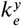) or rapid production and short pausing (high 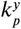 and 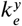). High production also implies that the proximal density of lysosomes will be higher, with approximately linear scaling (Figure 6B). We therefore used the measured ratio of LAMP1+ to LC3+ puncta in the proximal region to fit the appropriate value for 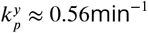.

Given the fitted model parameters, one key prediction is the spatial distribution of fused AVs. The fraction of AVs that have fused is plotted as a function of spatial position in Figure 6C, showing that much of the fusion occurs close to the distal tip, with only a gradual increase in fused fraction along the mid- and proximal axon. This result is consistent with experimental observations showing a relatively flat profile of fused fraction with distance from the distal tip (Figure 6C inset).

To further validate our mean-field model, we make use of stochastic simulations of lysosomal and AV particles in a linear domain (see Methods for details). The movement and switching of transport states for the simulated particles follow the same rules and use the same parameters as described above. Fusion occurs with probability *p*_*f*_ whenever the particles pass each other. Figure 6D shows a steady-state snapshot of particle states and positions from a simulation using the fitted parameters, and Figure 6–Supplemental Video 1 provides a corresponding video of the simulated particle motion. The spatial distributions of both LC3+ and LAMP1+ particles are similar for both the stochastic simulations and the mean-field model (Figure 6E). The simulations further enable direct tracking of where in the domain each individual AV becomes fused. This distribution of the position at first fusion (*x*_*f*_) shows a strong peak at the distal tip (Figure 6F). Given our fitted model parameters, we expect most AVs to fuse either in the distal tip or very soon after exit from the tip, with a smaller broad tail in the distribution corresponding to those that fuse throughout the rest of the axon. We note that this is a prediction of the model, which assumes fusion probability is constant regardless of position or motility state. The predominance of fusions in the far distal tip and their paucity in the mid- and proximal axon is a direct consequence of the observed organelle densities and motility patterns, and the observation that half of AVs in the distal axon are in a fused state. This prediction matches the observation that the fraction fused is largely invariant with distance from the tip, within the distal region of the axon (Figure 6C, inset).

Another prediction of the basic model described here is that the overall density of AVs must fall off with increasing distance from the origin, over a length scale of a few hundred micrometers. This is an inherent consequence of the assumption that AVs are produced in the distal tip and slowly transition to retrograde motility, with only 12% of the distal AVs observed to have made this switch. Because bidirectional organelles spread out slowly from their point of origin, these assumptions imply that the AVs must pile up in the distal region compared to elsewhere in the axon. A modest fall-off in the density of LC3+ puncta is indeed observed within the distal region (Figure 6E, inset). However, the observed LC3+ density is similar in both the distal and proximal regions (Figure 1D). This observation is at odds with the prediction of the model that the proximal AV density should be only 24% of the distal density. One potential explanation for this discrepancy is the non-linear geometry of the axon, with multiple distal tips potentially producing AVs that converge in the proximal region. We therefore expand our model to consider a branched axonal architecture.

#### Collateral branches supply AVs to maintain a broad axonal distribution

Neuronal axons form multiple branches known as axon collaterals. The generation of these collaterals enables a neuron to establish robust connectivity with neighboring targets and plays an important role in the development of the central nervous system (***Gallo, 2011***; ***Kalil and Dent, 2014***). We find that primary hippocampal neurons *in vitro* at DIV 7-10 have an average of *n*_*c*_ ≈ 5 axon collaterals per neuron with an average length of *L*_*c*_ ≈ 164μm (Figure 7—figure supplement 1). In order to represent this axonal geometry, we extend our linear mean-field model to include collaterals as one-dimensional branches growing at intervals from the main axon.

Our branched model geometry consists of a single main axon along with five collaterals placed at equispaced intervals along the mid-axon, excluding the most distal and most proximal 250μm regions (Figure 7A). Each collateral is taken to be a linear segment of equal length *L*_*c*_ ≈ 164μm. The end point of each collateral is assumed to be functionally equivalent to the main axon tip, producing AVs at a fixed rate 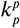 and allowing lysosomes to halt when they reach the distal tip, followed by returning at rate 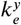 in the anterograde direction. Transport and fusion of AVs and endolysosomes along each branch and each segment of the main axon proceeds identically to the linear model. There are no preexisting studies on how anterograde organelles split between the main axon and a collateral when passing a junction point. In our model we make the relatively simple assumption that anterograde organelles split in proportion to the number of distal tips downstream of the junction. That is, the chances of entering the collateral are 1/2 at the most distal junction, 1/3 at the second-to-last junction, and so forth. This approach allows a similar number of organelles to reach each distal tip in the absence of fusion. Retrograde moving particles that pass a junction continue on upstream along the main axon.

**Figure 7.**
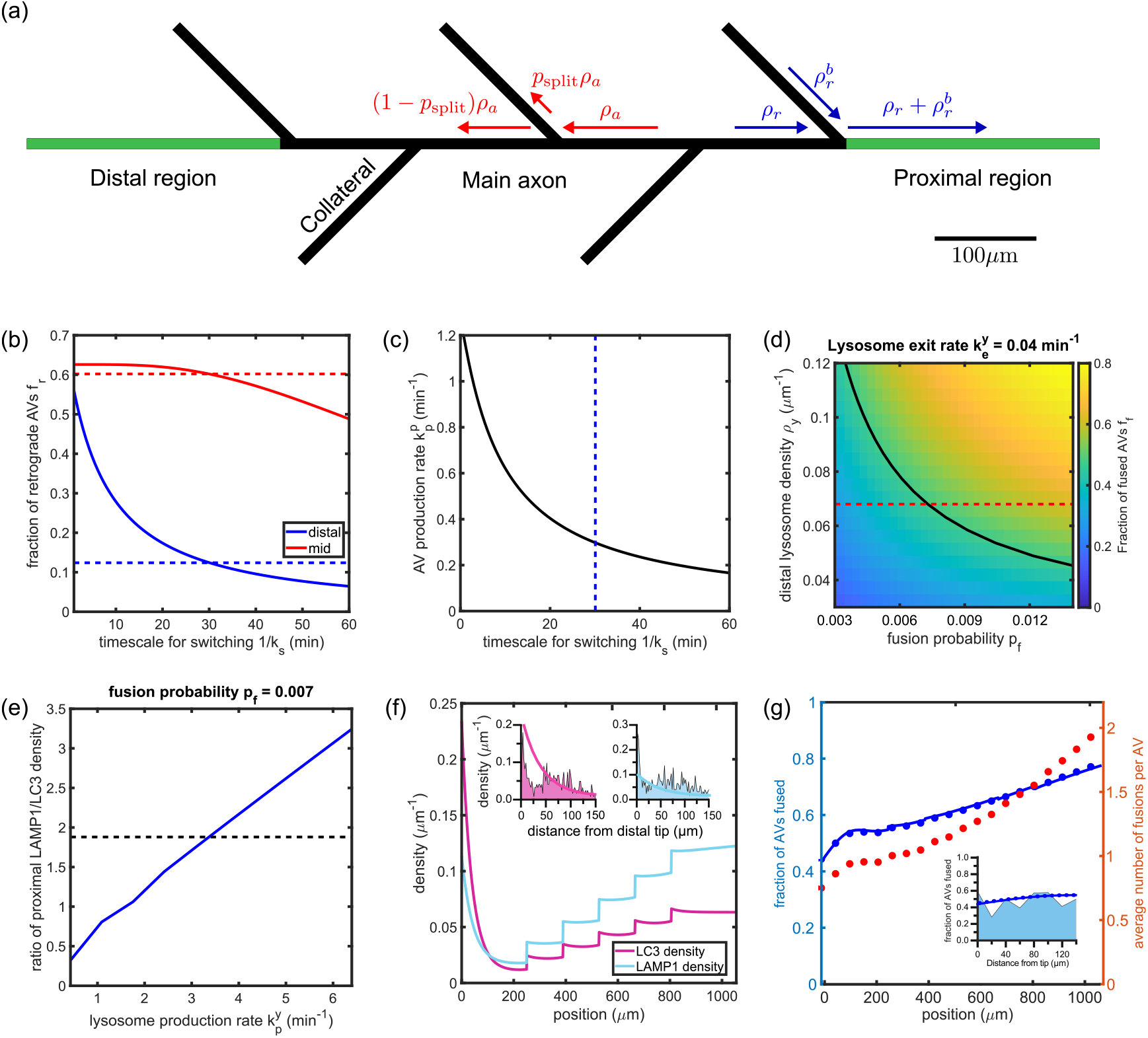
Mathematical model with branched axon morphology is consistent with measured organelle distributions. **(a)** Schematic of the branched axon geometry. All the distributions shown are within the main axon. Green sections indicate the extent of distal and proximal regions. Text in red and blue indicates the boundary conditions at branch junctions for anterograde and retrograde organelles, respectively. Scale bar: 100μm. **(b)** Fraction of AVs exhibiting retrograde motility within the distal (blue) and the mid (red) regions of the main axon, plotted against the timescale for switching. Observed fractions in hippocampal neurons are shown with corresponding dashed lines. **(c)** AV production rate 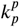 required to achieve the measured LC3+ density in the distal axon, plotted against the timescale for switching. The dashed line denotes the switching time obtained in **(b). (d)** Fraction of AVs fused within the distal axon *f*_*f*_, plotted against the fusion probability *p*_*f*_, and the lysosome density in the distal region. The tip-exit rate for lysosomes 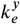 is set to 0.04 per minute. The solid black line denotes measured value of *f*_*f*_ based on LC3+ puncta colocalized with LAMP1+ puncta in the distal axon. The dashed red line denotes the density of LAMP1+ puncta observed in the distal axon. **(e)** The ratio of the lysosome density to AV density in the proximal axon, plotted against the lysosome production rate. Dashed black line denotes the measured value determined by enumerating LAMP1+ and LC3+ puncta in the proximal axon. **(f)** The linear density of LC3+ puncta (magenta) and LAMP1+ puncta (cyan) along the axon. The inset zooms into the distal region, showing the model prediction overlaid on experimentally observed LC3+ and LAMP1+ densities from Figure 1d and Figure 1f, respectively. **(g)** Spatial variation in the fraction of AVs fused at different positions along the axon. The solid blue line denotes the fraction fused in the base “one-and-done” model; blue markers give corresponding results for a modified model with unlimited fusion events (see Figure 7—figure supplement 2). The inset zooms into the distal region, overlaid with the experimentally observed distribution obtained by enumerating LC3+ puncta colocalized with LAMP1. Red markers denote the average number of fusions per AV for the unlimited fusion model. The following figure supplements are available for Figure 7: **Figure 7–Figure supplement 1**. Number and length of axon collaterals. **Figure 7–Figure supplement 2**. Parameter fitting for modified model with unrestricted fusion

The mean-field densities for AVs and lysosomes obey the same steady-state equations as the linear model (Eq. 1– 4), with boundary conditions at the junctions set according to the anterograde splitting law and the conservation of flux for retrograde organelles. The AV production rate 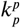, retrograde switch rate *k*_*s*_, and pausing rate for retrograde AVs 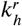 can again be obtained independently of the lysosome dynamics by fitting to experimental values of the distal AV density and the fraction of retrograde AVs in the distal and mid-axonal regions (Figure 7B,C). These fitted values to not differ substantially from the unbranched case. Parameters for fusion probability (*p*_*f*_) and lysosome dynamics 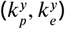 can also be obtained using similar methods as described for the linear model (Figure 7D,E). The fitted lysosome production rate for the branched model 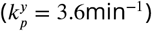 is roughly 6-fold higher than the linear model 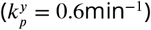 to enable a similar density of lysosomes to reach each individual distal tip. The fitted parameter values for the branched model are listed in Table 1.

The predicted spatial densities of LC3 and LAMP1 puncta along the main axon are shown in Figure 7F. At each branch junction point, the density of AVs increases as the retrograde organelles coming from the collateral join those moving along the main axon. The predicted average density of AVs in the proximal region is now approximately 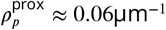, slightly higher than the distal density, and comparable to experimental measurements (Figure 1C,D). The model with branched axon geometry also yields a relatively flat profile for the fraction of AVs that have fused with a lysosome (Figure 7G). While the distal fraction fused 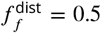 was used to fit the model parameters, the proximal fraction fused 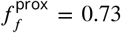 is a prediction of the model that approximately matches experimental measurements 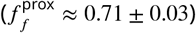 (Figure 3B).

#### Modeling unlimited fusion yields a relatively low number of fusion events

Parameterized off experimental measurements, the branched axon model of AV and lysosome dynamics accurately represents organelle densities in the distal and proximal regions, the typical fraction of fused organelles within each region, and the fraction of organelles in different motility states. We note that this model was developed under the assumption that each individual AV can fuse with at most one endolysosome. However, the actual number of fusion events between an autophagosome and lysosomes during transit along the axon is unknown. A variety of models could be developed wherein the probability of fusion *p*_*f*_ decreases as a function of the number of fusions already undergone or over time following the initial fusion event. The “one-and-done” model proposed here constitutes an extreme case where this decrease is very steep so that each AV immediately becomes incapable of fusion after the first such event.

We also consider a model for the opposite limit, where the number of fusions is unlimited and the fusion probability remains constant, regardless of how many previous endolysosomes have fused into a given AV. The equations for this alternate model are provided in the Methods section, and the resulting organelle densities and fraction fused profile are shown in Figure 7 – figure supplement 2. In principle, such a model could allow for a “snow-ball” effect where a single AV sweeps up large numbers of endolysosomes in successive fusions, leaving very few of them to reach the distal tips. However, given the fitted model parameters, we find that the average number of fusions accumulated by each AV is quite small: less than 1 in the distal axon, and rising to 2 fusions by the time an AV reaches the soma (Figure 7G, dashed red curve). This is a direct consequence of the low value of the fitted fusion probability *p*_*f*_ ≈ 0.007, which also leads to the fraction of AV with at least one fusion being very similar in both the “one-and-done” and the “unlimited fusions” model (Figure 7G,blue curves). Thus, a typical AV will have passed an average of ≈ 270 lysosomes by the time it reaches the soma, but will only have fused with a couple of them.

The available data described here does not allow us to distinguish whether or not there is a regulatory process that explicitly prevents an AV from fusing with multiple lysosomes. However, our quantitative model demonstrates that there is not a ‘snow-ball’ effect wherein individual AVs sweep up large numbers of lysosomes in multiple fusion events. Instead, the observed organelle distributions imply that the average number of fusions per AV is quite low, with only a small fraction of AV–endolysosome passage events resulting in fusion.

### Autophagosome Maturation in Axons is a Two-Step Process

Through fusion with an endolysosome, an AV acquires degradative hydrolases and a vATPase pump responsible for establishing and maintaining the acidic pH necessary for hydrolase function. In live-imaging studies, AV maturation can be assayed by measuring colocalization between fluorescent LC3 and the dye LysoTracker, which labels acidified compartments. In our primary hippocampal neurons, we find that the fraction of LC3+ puncta that colabel with LysoTracker is about 60% in the distal axon, with a slow increase to about 90% in the proximal axon (Figure 8A), similar to the early peak in fusion events and long tail observed in the modeling (Figure 6C,F). These observations are consistent with published data in multiple neuronal cell types examining the localization between LC3 and LysoTracker, LAMP1, and the late endosomal membrane protein Rab7 (Figure 8A, Table 2) (***Maday et al., 2012***; ***Maday and Holzbaur, 2014***; ***Cheng et al., 2015***; ***Cason et al., 2021***). Additionally, previous work using the fluorogenic enzymatic activity sensors MagicRed and MDW941, which specifically fluoresce when cleaved by the lysosomal hydrolases Cathepsin B and glucocerebrosidase respectively, showed similar high colocalization with LC3 in the distal axon (***Farfel-Becker et al., 2019***), again suggesting early fusion with active endolysosomes (Figure 8A, Table 2).

**Figure 8.**
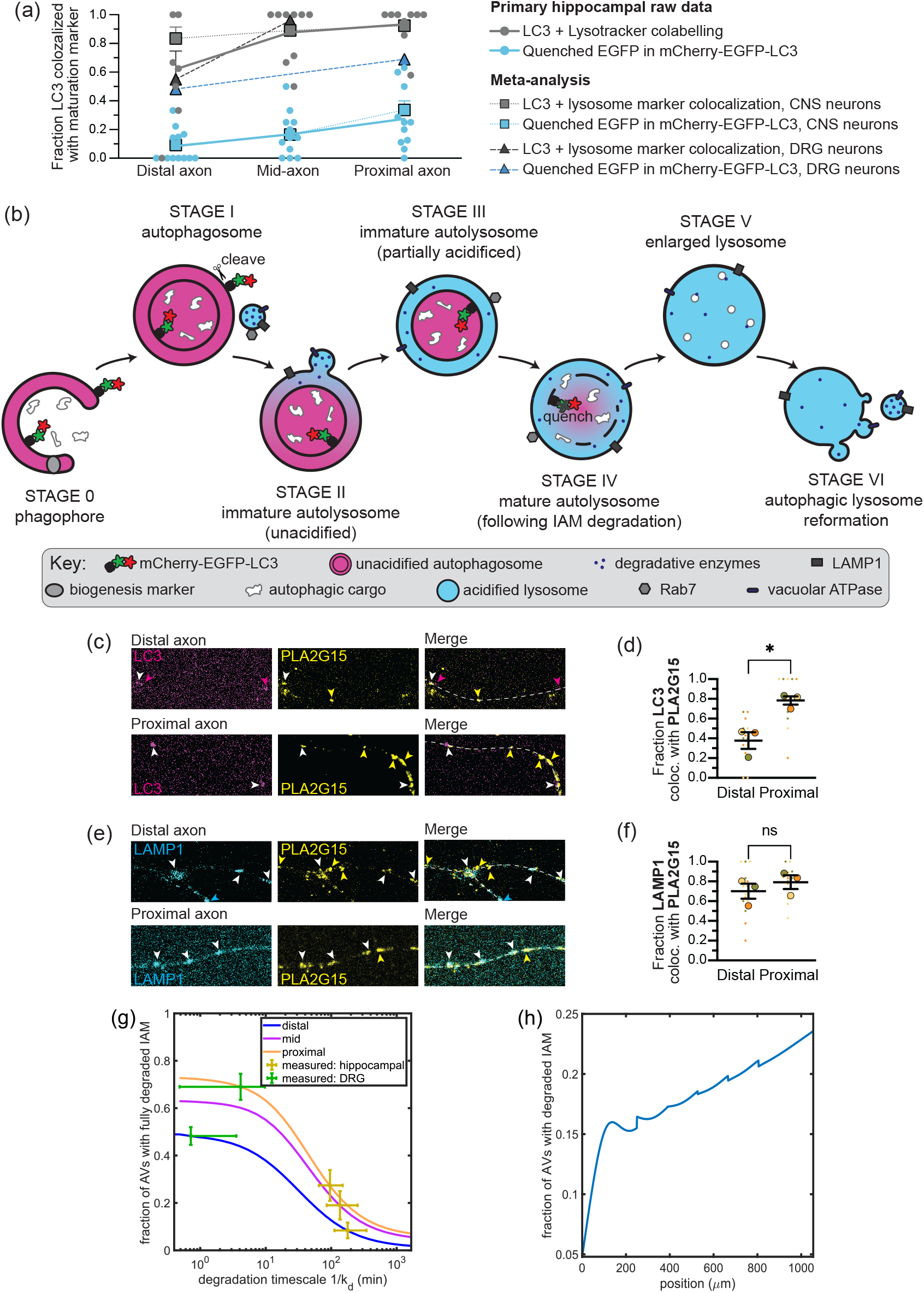
Distal fusion with endolysosomes is followed by slow inner membrane degradation. **(a)** Across studies and cell types, markers for AV-endolysosome fusion (LAMP1, Rab7, LysoTracker, MagicRed, MDW941) are acquired more distally than the GFP moeity of mCherry-GFP-LC3 quenches. Circular points represent primary hippocampal raw data, *n* = 8-12 axons. Lines represent quantities estimated from previous studies (see Table 2). **(b)** When an AV fuses with an endolysosome, the endolysosome’s contents enter the intermembrane space and its membrane proteins join the outer membrane. Following IAM degradation, the autophagic cargo, including mCherry-GFP-LC3, is exposed to lysosomal pH and proteases. Following degradation, the recyclable contents are exported and the remaining organelle is broken into smaller lysosomes. The rate of transition from fusion to membrane breakdown in denoted by *k*_*d*_. **(c)** Maximum projections showing LC3 and phospholipase A2 group XV (PLA2G15) overlap in the distal and proximal axon. **(d)** Comparison of the total LC3+ puncta colocalized with PLA2G15 in the distal and proximal axon. *n* = 3 trials; unpaired t test (*p* = 0.0127). **(e)** Maximum projections showing LAMP1 and PLA2G15 overlap in the distal and proximal axon. **(f)** Comparison of the total LAMP1+ puncta colocalized with PLA2G15 in the distal and proximal axon. *n* = 3 trials; unpaired t test (*p* = 0.4248). Magenta arrows, LC3 alone; yellow arrows, PLA2G15 alone; cyan arrows, LAMP1 alone; white arrows, colocalization. Scale bar, 5 μm. Fractions are all over the total LC3+ or LAMP1+ puncta in that region. Bars throughout show median ± 95% confidence interval. ns, p > 0.05; ***, p < 0.001. **(g)** Modeled fraction of AVs with degraded inner membrane *f*_*d*_, plotted as a function of the degradation time (*τ*_*d*_ = 1/*k*_*d*_). Plots shown are averages over the most distal (blue) and most proximal (orange) 250μm regions, along with the middle section (pink) of a modeled axon of length 1055μm. Parameters used in the model are the same ones extracted from Figure 7. The degradation times corresponding to measured values of *f*_*d*_ in the distal, mid-axon, and proximal regions are shown in yellow for hippocampal neurons, and in green for DRGs. **(h)** Distribution of the fraction of AVs with degraded IAM along the axon. The degradation rate is chosen to be the average of predicted values from **(g)** for hippocampal neurons.

**Table 2.** Meta-analysis of autophagosome maturation in axons

| Cell type | Marker | Distal | Mid | Proximal | Reference |
| --- | --- | --- | --- | --- | --- |
| Hippocampal | Rab7 | 88% | 88% | 85% | *Cason et al. (2021)* |
| Hippocampal | LAMP1 | 84% | 91% | 99% | *Cason et al. (2021)* |
| Hippocampal | LysoTracker | 62% | 88% | 93% | *Cason et al. (2021)* |
| Hippocampal | quenched EGFP | 8% | 17% | 27% | *Cason et al. (2021)* |
| Cortical | MagicRed | 100% | – | – | *Farfel-Becker et al. (2019)* |
| iPSC-derived | quenched EGFP | 10% | – | 40% | *Boecker et al. (2021)* |
| DRG | Rab7 | 95% | 95% | – | *Cheng et al. (2015)* |
| DRG | LAMP1 | 10% | 97% | – | *Maday et al. (2012)* |
| DRG | LysoTracker | 35% | 96% | – | *Maday et al. (2012)* |
| DRG | MDW941 | 80% | – | – | *Farfel-Becker et al. (2019)* |
| DRG | quenched EGFP | 48% | – | 69% | *Maday et al. (2012)* |

However, use of the dual-color LC3 reporter mCherry-EGFP-LC3 shows a very different rate of AV maturation. This reporter fluoresces in the red and green wavelengths in nonacidified environments, but only red in acidic environments due to the quenching of the EGFP moiety below pH 5.8 (***Campbell and Choy, 2001***; ***Pankiv et al., 2007***). Thus, AVs fluorescing red and green can be considered “immature” while those showing only mCherry fluorescence are “mature.” Use of this reporter in primary hippocampal or iPSC-derived cortical neurons revealed much slower acidification than LysoTracker, with almost no AVs in the distal axon showing acidity-triggered EGFP quenching (Figure 8A, Table 2) (***Cason et al., 2021***; ***Boecker et al., 2021***). Furthermore, while all of the other lysosomal markers labeled the vast majority of AVs by the mid- or proximal axon, less than half of AVs in the proximal axon of primary hippocampal or iPSC-derived cortical neurons demonstrated EGFP quenching (***Cason et al., 2021***; ***Boecker et al., 2021***). The marked difference between EGFP quenching and AV colabeling with other lysosome markers in these central nervous system (CNS) neurons is illustrated in Figure 8A. Previous work in dorsal root ganglia (DRG) neurons likewise shows a delay in EGFP quenching as compared with other markers of lysosomal fusion, although the gap is not as large (Figure 8A, Table 2) (***Maday et al., 2012***).

The double-membraned nature of AVs is key to reconciling these seemingly disparate observations (Figure 8B). During biogenesis, the growing phagophore (Stage 0) engulfs cargo such that the cargo ends up inside the inner autophagosomal membrane (IAM), within the central lumen of the closed autophagosome (Stage I). When an autophagosome and an endolysosome fuse, the endolysosomal membrane becomes part of the outer membrane and the contents of the endolysosomal lumen are delivered into the intermembrane space. Thus, degradative enzymes and H^+^ ions from the endolysosome specifically occupy the space between the outer and inner membranes. Because the volume of the intermembrane space is larger than the volume of the endolysosome, fusion will cause the pH to rise transiently (Stage II); however, since the fraction of AV colocalized with endolysosomal markers (Figure 3) is similar to the fraction of LysoTracker+ AVs (Figure 55E), acidification of the intermembrane space, achieved by activity of vATPase pump(s) in the outer membrane, likely occurs rapidly following fusion (Stage III). In non-neuronal cells (mouse embryonic fibroblasts), LysoTracker has been shown to specifically localize to the intermembrane space and then collapse inward when the IAM is degraded (***Tsuboyama et al., 2016***).

Axonal AVs are condensed due to the narrow diameter of the axon and rarely appear as rings after leaving the tip, making this intermembrane space and inner membrane collapse difficult to resolve with conventional fluorescence microscopy. However, once the IAM is broken down we would expect the central lumen and the cargo therein to be exposed to the acidic pH required to activate enzymatic degradation (Stage IV). Unlike LysoTracker and the fluorogenic enzyme activity sensors, the tandem mCherry-EGFP-LC3 marker localizes specifically to the inner lumen of the AV (Figure 8B). Initially LC3 localizes to both the inner and outer membranes, conjugated to the lipid phosphatidylethanolamine so the protein extends into the lumen on the inner membrane and into the cytosol on the outer membrane (***Martens and Fracchiolla, 2020***). The protein extending into the cytosol is cleaved by the autophagy protease ATG4 (***Kauffman et al., 2018***) leaving only the lumenal protein, and fluorophores, intact. Therefore quenching of the EGFP moiety can be used as a specific readout of IAM degradation, indicating the point at which the IAM breaks down and the lumenal LC3 and other cargo are exposed to the acidic pH and degradative enzymes (Figure 8B).

A careful comparison of the tandem mCherry-EGFP-LC3 marker and other endolysosomal markers thus reveals autophagic maturation to be a two-step process. The first step involves fusion with one or more endolysosomes to acquire degradative enzymes and trigger acidification of the intermembrane space, yielding an immature autolysosome. The second step involves breakdown of the inner autophagosomal membrane to enable the enzymes and acidic environment to reach the lumen, yielding a mature autolysosome.

#### Modeling slow IAM degradation

Several explanations could account for the distinct spatial distributions of AV-lysosome fusion and IAM degradation in hippocampal neurons. One possibility is that the endolysosomes which fuse with AVs in the distal region may be lacking the enzymes responsible for IAM degradation. Recently the lysosomal lipase LPLA-2 was identified in *C. elegans* through a forward genetic screen as the enzyme responsible for IAM breakdown (***Li et al., 2021***). We probed for the mammalian ortholog phospholipase A2 group XV (PLA2G15) in fixed primary hippocampal neurons and found that endogenous PLA2G15 was present in axons and colocalized with both endogenous LC3 and LAMP1 (Figure 8C-F). Specifically, about 40% of LC3+ puncta in the distal axon colocalized with PLA2G15, increasing to about 80% in the proximal axon (Figure 8D). The majority of LAMP1+ puncta colocalized with PLA2G15 throughout the axon, with no significant difference between the distal and proximal axon (Figure 8F). The colocalization between PLA2G15 and both LC3 and LAMP1 was very similar to the colocalzation with other lysosomal enzymes and the vATPase (Figures 2-3). We therefore conclude that the phospholipase required for IAM degradation is acquired in the initial AV-endolysosome fusion event.

An alternative explanation for the different spatial profiles of AV-endolysosome fusion and IAM degradation is a temporal gap: slow kinetics of the IAM degradation could allow time for the AVs to reach the soma in many cases before completion. We explore this possibility, leveraging our parameterized one-and-done model for AV fusion in a branched axonal geometry (Figure 7). Namely, we introduce a single new parameter *k*_*d*_ describing a constant-rate process for complete inner membrane degradation in an AV that has undergone fusion with an endolysosome. The model is then expanded to include a new set of states and spatial distributions 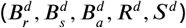 corre-sponding to bidirectional, processively retrograde, and paused AV densities where IAM degradation has been completed, following fusion with an endolysosome. The steady-state distributions of such organelles obey a set of inhomogeneous linear equations that are solved as described in Methods.

To obtain an estimate of the degradation rate *k*_*d*_, we consider the fraction of all AVs that are in the IAM-degraded state 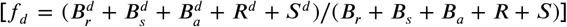 averaged over the distal and proximal axonal regions. These fractions are plotted as a function of the degradation timescale *τ*_*d*_ = 1/*k*_*d*_ in Figure 8G. A single value of *τ*_*d*_ ≈ 100min can account for the experimentally observed low values of degraded fraction in both the distal 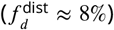 and proximal 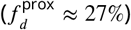 axon of hippocampal neurons. This timescale can be compared to the average time required for a newly formed AV to enter the processive retrograde state (1/*k*_*s*_ ≈ 30min) and to move from the distal tip to the soma (*L*/*v*_*p*_ ≈ 20min) in the fitted model. The predicted spatial profile for AVs with a degraded inner membrane is nearly flat throughout most of the axon (Figure 8H), analogous to the fraction of AVs that have undergone a fusion event.

We find it takes ≈ 100min following fusion to degrade the IAM of AVs in primary hippocampal neurons (Figure 8G). The slow degradation kinetics account for the relatively small fraction of AVs whose IAM is fully degraded by the time they reach the soma. By comparison, the larger fraction of quenched EGFP in both proximal and distal regions of DRG neurons (Table 2) implies a much shorter IAM degradation timescale in these neurons, similar to that seen in mouse embryonic fibroblasts (***Tsuboyama et al., 2016***). Although detailed fitting is not carried out for this alternate cell type, using the parameters obtained for hippocampal cells yields a rough estimate of *τ*_*d*_ ≈ 1− 10 min for the IAM degradation time in DRG neurons (Figure 8G). This could perhaps be due to higher expression levels of phospholipase and/or enhanced phospholipase activity, possibilities to be explored in future studies.

## Discussion

The results of this study provide a quantitative perspective on the maturation of neuronal autophagosomes, a key aspect of the autophagy pathway crucial to maintaining the recycling and turnover of cell components. Previous work both *in vitro* and *in vivo* showed that autophagic vacuoles (AVs) form within the distal axon, then fuse with endolysosomes while moving in a retrograde fashion towards the soma (***Maday et al., 2012***; ***Stavoe et al., 2016***; ***Hill et al., 2019***). Fusion with endolysosomes provides AVs with degradative enzymes necessary to break down their cargo, as well as the vATPase pump that is necessary to establish and maintain the acidic pH at which degradative enzymes are active (***Yin et al., 2016***) However, the location and timing of fusion events and their relationship to AV maturation remained largely unclear.

We leveraged endogenous staining of AV and endolysosomal proteins to quantify the colocalization and spatial distribution of these organelles along the axons of primary rat hippocampal neurons, thereby avoiding potential overexpression artifacts. We find that LAMP1 colocalizes readily with both degradative hydrolases and active vATPase in both the distal and proximal axon (Figure 2). Note that our immunofluorescence is not quantitative and we are unable to detect the number of vATPase complexes present on each endolysosomal structure. Recent work in nonneuronal cells would predict endolysosomes in the periphery of the cell likely have only one vATPase each (***Maxson et al., 2022***). We also cannot quantify the relative load of degradative enzymes, and our CTSL antibody weakly cross-reacts with the immature form of CTSL, pro-CTSL; however, the presence of AEP, a protease known to cleave pro-CTSL into mature CTSL, suggests mature CTSL is the predominant form (***Maehr et al., 2005***). Thus, while we cannot measure degradative activity in a fixed assay, we conclude the LAMP1+ endolysosomes in the axon are degradatively competent.

We find that roughly half of AVs have fused with at least one competent endolysosome by the time they leave the distal axonal region, as measured by colocalization with LAMP1, degradative enzymes, and the vATPase (Figure 3). Live-cell imaging of fluorescently tagged LC3 was used to establish the motility behavior of axonal AVs (Figure 5). We find that the majority of AVs in the distal axon are stationary or display short, unbiased bidirectional motions, and that the majority of AVs in the mid- and proximal axon move retrograde towards the soma, consistent with previous studies (***Maday et al., 2012***; ***Maday and Holzbaur, 2014***; ***Cheng et al., 2015***; ***Cason et al., 2021***; ***Boecker et al., 2021***). Furthermore, we find that endolysosomal fusion is independent from initiation of retrograde transport, with no difference in motility measurements between AVs positive or negative for the endolysosomal marker LysoTracker (Figure 5E,F).

While direct measurements of colocalization between different organelle labels can indicate whether at least one fusion event has occurred, they cannot establish how many endolysosomes a single AV has fused with, nor how many endolysosomes have passed without fusion. We therefore developed a mean-field mathematical model to describe the motility and fusion interactions between AVs and endolysosomes, which allows us to translate the experimental measurements into a quantitative picture of the behavior of these organelles. Our model is parameterized in such a way as to reproduce a variety of experimental metrics, including distal and proximal densities of AVs and endolysosomes, the fraction of fused AVs in the distal region, and the fraction of retrograde-moving AVs. The model shows that fusion of an AV with an endolysosome is expected to be a rare event, with fewer than 1% of passage events resulting in a fusion. It also predicts that a large fraction of AVs will undergo their first fusion while still in the distal region of the axon, with only a gradual increase in fusions thereafter (Figure 6), consistent with observations in fixed neurons.

Notably, we find that each AV is expected to fuse with only one to a few endolysosomes by the time it reaches the soma. The comparison between quantitative modeling and experimental data rules out the possibility of ‘snow-balling’ AVs that soak up large numbers of endolysosomes while moving through the axon. This effect could be achieved by regulatory mechanisms that restrict subsequent fusions (***Saleeb et al., 2019***) or simply as a result of the low probability of fusion upon passing (***Saleeb et al., 2019***; ***Li et al., 2020***; ***Shen et al., 2021***). Specifically, we hypothesize that the regulation of the SNARE protein syntaxin-17 (Stx17) may account for the relatively low number of fusion events predicted by the models. Stx17 resides in the AV outer membrane and forms a complex with the SNAREs synaptosome-associated protein 29 (SNAP29) and vesicle-associated membrane proteins 7 and 8 (VAMP7/8) to facilitate fusion(***Itakura et al., 2012***). Stx17 is tightly regulated via posttranslational modifications, autoinhibition, and interaction with lysosomal membrane proteins to prevent ectopic fusion events (***Saleeb et al., 2019***; ***Li et al., 2020***; ***Shen et al., 2021***). Regardless of the precise mechanism, the limited number of fusions allows for a broad distribution of endolysosomes throughout the axon, making them available for interaction with other organelles even in regions far away from the proximal axon where they are produced.

An interesting feature that arises from the quantitative model is the importance of axon geometry in modulating the interactions and distribution of organelles. Specifically, a linear geometry with AVs produced at the distal tip results in accumulation of AVs in the distal region and their depletion in the proximal axon. This is a direct consequence of the initial bidirectional motion near their production site, followed by unidirectional retrograde transport to an absorbing boundary at the soma. However, in a branched axon where AVs are also produced at collateral branch tips, the AVs are expected to be more broadly distributed throughout the axon (Figure 7). The lysosome production rate at the soma must be ramped up concomitantly to enable a sufficient density of lysosomes to reach the main axon and collateral branch distal tips.

Fusion with an endolysosome is only the first step in the maturation of an AV. Our results show that nearly half of axonal AVs fuse with an endolysosome in the distal axon and exhibit concomitant partial acidification as marked by the pH-sensitive LysoTracker dye. However, only a small fraction of AVs become fully acidified, as indicated by the quenching of the EGFP moiety of mCherry-EGFP-LC3 in the AV lumen, by the time they reach the proximal axon (Figure 8A). These observations support a two-step model of autophagosome maturation, wherein fusion with an endolysosome allows for the acquisition of endolysosomal markers and acidification of the space between the outer and inner autophagosomal membrane (IAM), followed by the relatively slow degradation of the IAM (***Tsuboyama et al., 2016***). It is only when the IAM is degraded that the AV lumen, including mCherry-EGFP-LC3 and the autophagic cargo, becomes fully acidified and cargo degradation may begin (Figure 8B). This model is consistent with data from multiple neuronal cell types, wherein acquisition of endolysosomal markers precedes quenching of the mCherry-EGFP-LC3 reporter (Table 2, Figure 8A).

We incorporate the additional IAM degradation step into our model, and extract a quantitative estimate of the rate for this process (Figure 8I) using mCherry-EGFP-LC3 quenching as a readout of IAM degradation. Notably, mCherry-EGFP-LC3 quenching is a marker for IAM degradation only if the mCherry-EGFP-LC3 proteins on the outer autophagosomal membrane, which extend into the cytosol, are cleaved. The protease ATG4 is responsible for cleaving LC3 and other members of its protein family from the outer autophagosomal membrane (***Kauffman et al., 2018***). Given that ATG4s are also involved in autophagosome formation (***Fujita et al., 2008***; ***Agrotis et al., 2019***), they are likely to be enriched in the distal axon and therefore we anticipate LC3 is rapidly cleaved from nascent autophagosomes in the distal axon. Furthermore, work in *C. elegans* neurons showed that ATG4 activity was required for autophagosome-lysosome fusion (***Hill et al., 2019***), thus mCherry-EGFP-LC3 should be removed from the outer autophagosomal membrane prior to fusion with endolysosomes. Therefore we conclude mCherry-EGFP-LC3 quenching is more likely a readout of IAM degradation rather than a readout of ATG4 activity.

We find that a single IAM degradation rate constant is consistent with the mCherry-EGFP-LC3 quenching measurements taken in both the distal and the proximal axon regions. The average IAM degradation time in primary hippocampal neurons (*τ*_*d*_ ≈ 100min) is more than an order of magnitude longer than the time between fusion and IAM breakdown observed in mouse embryonic fibroblasts (≈6.6min) (***Tsuboyama et al., 2016***). However, the rate extracted for DRG neurons (Figure 8G), derived from previously published data (***Maday and Holzbaur, 2014***), is relatively similar to that seen in mouse embryonic fibroblasts (***Tsuboyama et al., 2016***). This highlights a difference not only between neurons and non-neuronal cells, but also between neuronal cell types that will need to be reconciled by future experimentation.

The modeling approach developed here serves as a framework for quantitatively understanding how the interplay between organelle transport and interactions across space and time governs autophagosome maturation. By combining modeling with direct measurements of organelle motility in live neurons and fusion under endogenous conditions we have reconciled multiple conflicting studies of AV maturation and quantitatively connected organellar transport and fusion. Because autophagy defects are implicated in a variety of neurodegenerative diseases, obtaining a clear picture of this pathway is an important step towards a mechanistic understanding of such disorders.

## Materials and Methods

### Primary hippocampal culture

Sprague Dawley rat hippocampal neurons at embryonic day 18 were obtained from the Neurons R Us Culture Service Center at the University of Pennsylvania. Cells (immunofluorescence, 180,000 cells; live imaging, 200,000 cells) were plated in 20 mm glass-bottom 35 mm dishes (MatTek) that were precoated with 0.5 mg/ml poly-L-lysine (Sigma Aldrich). Cells were initially plated in Attachment Media (MEM supplemented with 10% horse serum, 33 mM D-glucose, and 1 mM sodium pyruvate) which was replaced with Maintenance Media (Neurobasal [Gibco] supplemented with 33 mM D-glucose, 2 mM GlutaMAX (Invitrogen), 100 units/ml penicillin, 100 mg/ml streptomycin, and 2% B-27 [ThermoFisher]) after 5-20 h. Neurons were maintained at 37 C in a 5% CO2 incubator; cytosine arabinoside (Ara-C; final conc. 1 μM) was added the day after plating to prevent glia cell proliferation. Where applicable, neurons (5-7 DIV) were transfected with 0.35–1.5 μg of total plasmid DNA using Lipofectamine 2000 Transfection Reagent (ThermoFisher, 11668030) and incubated for 18-24 h.

### iPSC-derived neuron culture

Induced pluripotent stem cells (iPSC) from the KOLF2.1J lineage were cultured, induced, and transfected exactly as described in ***Pantazis et al***. (***2021***) with the following exception: to stably express doxycycline-inducible hNGN2 using a PiggyBac delivery system, iPSCs were transfected with PB-TO-hNGN2 vector (gift from M. Ward, NIH, Maryland) in a 1:2 ratio (transposase:vector) using Lipofectamine Stem (ThermoFisher); after 72 hours, transfected iPSCs were selected for 48 hours with 0.5 μg/mL puromycin (Takara).

### Immunofluorescence experiments and analysis

Neurons were fixed at 7-10 days *in vitro* for 30 minutes at room temperature using Bouin’s solution (SigmaAldrich, HT10132) supplemented with 8% sucrose and diluted 50% in Maintenance Media. Bouin’s solution was then removed and the cells were washed in PBS before being stored for up to 6 months in PBS at 4°C. Cells were permeabilized for 8 minutes at -20ºC in Optima Methanol (ThermoFisher, A456-1)and washed in PBS, then blocked for 1 hour at room temperature in blocking solution (5% normal goat serum, 1% bovine serum albumin, 0.05% sodium azide). Primary and secondary antibodies (see Table 3 for manufacturers and dilutions) were diluted in blocking solution and each left on cells for 1 hour at room temperature, with 3×5 min washes in PBS after each incubation. Cells were mounted in Prolong Gold (ThermoFisher, P36930) and imaged within 48 hours at 100x on a Perkin Elmer UltraView Vox spinning disk confocal on a Nikon Eclipse Ti Microscope with a Plan Apochromat Lambda 60x 1.40 NA oil-immersion objective and a Hamamatsu EMCCD C9100-50 camera driven by Volocity (PerkinElmer). Z stacks were acquired in 0.1 − 0.2μm steps.

**Table 3.** Reagents used in the study

| Primary antibodies |  |  |  |  |  |
| --- | --- | --- | --- | --- | --- |
| Target | Host | Dilution | Manufacturer | Cat# | RRID |
| LC3 | Rabbit | 1:250 | Abcam | ab48394 | AB_881433 |
| LC3 | Mouse | 1:50 | Santa Cruz | sc-376404 | AB_11150489 |
| LAMP1 | Sheep | 1:50-100 | R and D Systems | AF4800 | AB_1026176 |
| LAMP1 | Rat | 1:50-100 | DSHB | 1d4b | AB_2134500 |
| AEP | Sheep | 1:100 | R and D Systems | AF2058 | AB_2234536 |
| CTSL | Mouse | 1:100 | Novus | NB100-1775 | AB_10124480 |
| ATP6V1F | Mouse | 1:100 | Novus | NBP2-03498 | AB_2904246 |
| PLA2G15 | Rabbit | 1:125-150 | Biorbyt | orb185108 | AB_2904247 |
| Tau | Chicken | 1:300 | Synaptic Systems | 314 006 | AB_2620049 |

| Secondary antibodies |  |  |  |  |
| --- | --- | --- | --- | --- |
| Target | Conjugation | Dilution | Manufacturer | Cat# |
| Sheep | Alexa Fluor 405 | 1:1000 | Abcam | ab175676 |
| Rat | Alexa Fluor 405 | 1:1000 | Abcam | ab175671 |
| Chicken | Alexa Fluor 488 | 1:1000 | ThermoFisher | A11039 |
| Rabbit | Alexa Fluor 555 | 1:1000 | ThermoFisher | A21429 |
| Mouse | Alexa Fluor 555 | 1:1000 | ThermoFisher | A21424 |
| Sheep | Alexa Fluor 594 | 1:1000 | ThermoFisher | A11016 |
| Rabbit | Alexa Fluor 647 | 1:1000 | ThermoFisher | A31573 |
| Mouse | Alexa Fluor 647 | 1:1000 | ThermoFisher | A32728 |

| Materials for live-cell imaging |  |
| --- | --- |
| Material | Source |
| mCherry-EGFP-LC3 | Gift from T. Johansen, University of Tromsø |
| mScarlet-LC3B | Subcloned from Addgene #21073 and Addgene #85054 |
| LAMP1-mNeon | Subcloned from Addgene #98882 into PGK vector |
| LysoTracker DeepRed | ThermoFisher Cat #L12492 |

Analysis was performed on maximum z projections in ImageJ (https://imagej.net/ImageJ2). Using the tau staining, axons were straightened (line width = 20 pixels) specifically in regions where they did not overlap with other cells. Because of the potential for cytoplasmic background staining, analysis was performed manually. However, to avoid bias, any immunofluorescence under an specific intensity threshold (0.5% for LC3 antibodies; 1% for all other antibodies) was excluded. LC3+ or LAMP1+ structures were defined as punctae ≥ 2 and < 20 pixels in diameter and signals within 7 pixels (≈ 1μm) were considered colocalized.

### Live-cell neuron imaging and analysis

Where applicable, neurons were incubated with LysoTracker (25 nM) for 15-30 min, which was then removed for imaging. Neurons were imaged in Imaging Media (HibernateE [Brain Bits] supplemented with 2% B27 and 33 mM D-glucose). Autophagosome behavior was monitored in the proximal (<250 μm from the soma), distal (<250 μm from the distal tip), or mid-axon of 7–8 DIV neurons imaged at a rate of 1 timepoints/sec for 1-3 min. Neurons were imaged in an environmental chamber at 37ºC with a Apochromat 100 × 1.49 numerical aperture (NA) oil-immersion objective on the spinning disk confocal described above. Only cells expressing moderate levels of fluorescent proteins were imaged to avoid overexpression artifacts or aggregation. It should be noted that the quality of the primary neuron dissections can affect autophagosomal motility, leading to variable retrograde fractions.

Kymographs were generated in ImageJ using the MultiKymograph plugin (line width = 5) and analyzed either in ImageJ. Puncta were classified as either anterograde (moving ≥10μm towards the axon tip), retrograde (moving ≥10μm towards the soma), or stationary/bidirectional (net movement ≤10μm during the video). Because fluorescent LC3 is cytosolic (as well as punctate) and neurites occasionally crossed in culture, raw videos were referenced throughout kymograph analysis to ensure only real puncta (≥ 1.5 SD from the axon mean) were included in analyses. All comigration analyses were performed using kymographs.

### Statistics for cell-based experiments

All statistical analyses were performed in Prism (GraphPad, San Diego, CA). Unless otherwise indicated, n indicates the number of trials (superplotting) wherein at least 3 cells were analyzed per trial. Neither parametricity nor preemptive sample-size (power) analyses were performed; however data appears normally distributed and post-hoc power calculations were used to confirm a sufficient number of replicates were collected. Statistical measures are described in the legends.

### Parameter estimates for bidirectional motility

We use live-cell dynamic imaging to extract estimates of the parameters describing AV bidirectional motility. Kymographs for LC3+ puncta within the distal 250*μ*m of the axon were obtained at a temporal resolution of 1frame/sec, for a total imaging period of 1 − 3 min. Manual tracing was used to extract a total of 49 AV trajectories from the kymographs. For these trajectories, the net displacement was used to classify AVs undergoing long-range retrograde motion (> 10*μ*m towards the soma) or anterograde motion (> 10*μ*m towards the tip), with the remaining particles classified as in a bidirectional/stationary state. Among the bidirectional/stationary particles, those whose trajectory showed a range (maximal minus minimal position) below 3*μ*m were classified as stationary and the rest as bidirectional. Among the bidirectional trajectories, we extracted all segments where the particles moved in a consistent direction (anterograde or retrograde) and found that the average displacement during such segments was *λ* ≈ 1.82± 0.16*μ*m, motivating our choice of a 2*μ*m run-length in the model.

### Steady state solutions for basic mathematical model

The steady-state densities of AVs in different motility states on a linear domain (Eq. 1) were solved using elementary matrix methods for a set of homogeneous first-order differential equations with constant coefficients ***Boyce et al***. (***2021***). To solve for the steady-state distributions of unfused AVs and endolysosomes (Eq. 2–4), we used the built-in solver `bvp4c` in Matlab ***MATLAB*** (***2021***), which provides a 4th-order method for solving boundary value problems on a set of linear regions. Code for implementing the model for a given set of parameters is provided at https://github.com/lenafabr/autophagyTransportModel.

The process of solving for state densities on a branched axon remains the same as that for the linear model within each branch and each contiguous segment of the main axon. Additional boundary conditions at the junctions are given by:

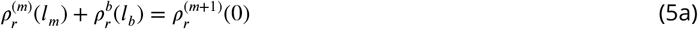

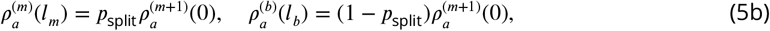

where 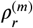 and 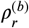 denote densities of retrograde organelles (*B*_*r*_, *R, Y*_*r*_) on a main axon segment and on a branch, respectively; 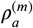 and 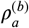 denote densities of anterograde organelles (*B*_*a*_, *Y*_*a*_), and *l*_*m*_, *l*_*b*_ are the length of the corresponding main segment and branch. The splitting of densities at each junction is determined by *p*_split_ = 1/(*b* + 1), which is defined such that the fraction of organelles proceeding each main segment is proportional to the number of distal tips downstream of that segment.

### Modified model with unlimited fusions

For the alternate model where each AV can fuse with an unlimited number of endolysosomes, Eq. 1 for the total AV density in different motility states, and Eq. 2 for the densities of unfused AVs, remain valid. The densities of endolysosomes are described by the following equations:

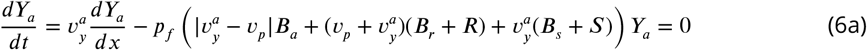

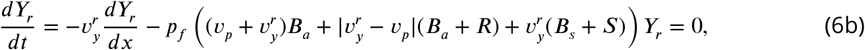

which enable fusion with all AVs regardless of their prior fusion state. The corresponding boundary conditions are identical to Eq. 4b-c, with an altered condition on endolysosomes at the distal tip:

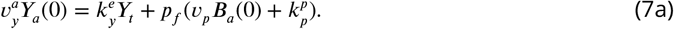

For this set of equations, the endolysosome densities are computed by direct integration. The densities of unfused AVs (Eq. 2, 4b,c) are found using the boundary-value problem solver `bvp4c`.

An additional metric of interest for this model is the average number of fusions undergone by AVs found in different regions of the axon. We define the densities 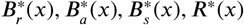 to correspond to the linear density of fusion counts in AVs that are in each of the motility states. For example, 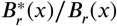 gives the average number of fusions among the bidirectional retrograde AVs found at position *x* along the axon. These fusion densities obey the following set of steady-state equations:

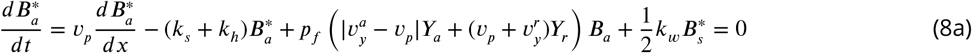

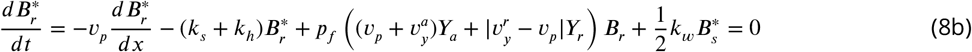

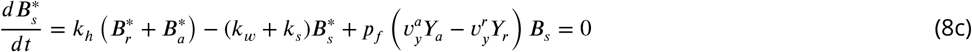

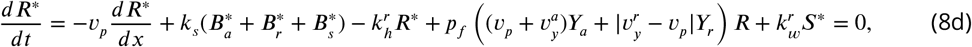

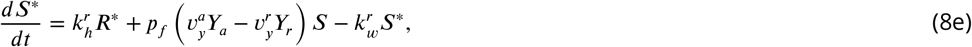

with boundary conditions at the distal tips and the soma:

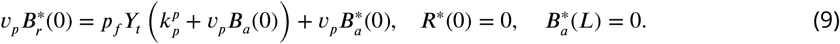

Once the endolysosome densities are computed, Eq. 8– 9 form a set of linear nonhomogeneous equations that are solved using standard matrix methods (***Boyce et al., 2021***).

### Steady state solutions for IAM degradation model

We define the density 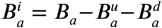, for AVs in the bidirectional anterograde state that have fused with an endolysosome but have not yet undergone full IAM degradation. Analogous densities are defined for the other motility states 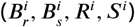

The dynamic equations for these fused AVs with intact IAM at steady state are given by

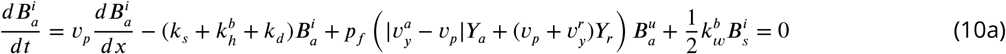

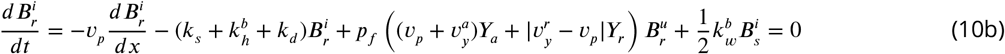

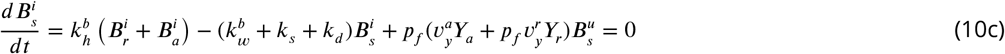

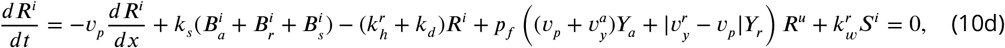

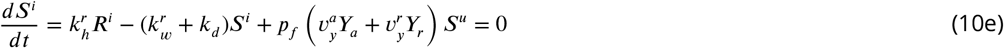

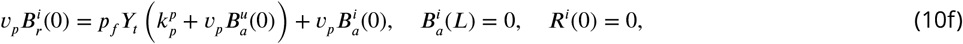

where *k*_*d*_ denotes the degradation rate and the endolysosome densities *Y*_*a*_, *Y*_*r*_, as well as the unfused AV densities 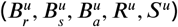 are precalculated as previously described. Equations 10 can then be treated as a system of nonhomogeneous linear equations, solvable via standard matrix methods (***Boyce et al., 2021***).

### Agent-based stochastic simulations for interacting organelles

Organelle interactions are simulated explicitly using custom written FORTRAN 90 code available at: https://github.com/lenafabr/particleDynamics1D. We simulate a linear domain (0 < *x* < *L*) of length *L* = 1055μm with *x* = 0 denoting the distal axonal tip, and *x* = *L* representing the soma. Point-particle endolysosomes spawn at the soma (*x* = *L*) and move towards the distal tip with a velocity 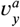. Upon reaching the distal end, the endolysosomes halt at the tip before engaging in retrograde motility at a rate 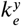 with a velocity 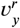. Point-particle AVs spawn at the distal tip (*x* = 0) in a bidirectional state moving in the retrograde direction. Motile AVs in the bidirectional state can halt with a rate *k*_*h*_. Halted AVs resume motion at rate *k*_*w*_, equally likely in the retrograde or anterograde direction. AVs from all bidirectional states can switch to processive retrograde state at a rate *k*_*s*_. All motile AVs move at a velocity *v*_*p*_, and have a rate 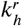 for pausing and a rate 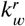 for restarting motility.

Interactions between an endolysosome and an AV occur whenever the two organelles cross past each other. Each such passage event results in fusion with a probability *p*_*f*_. A fusion event destroys the lysosome, while the AV particle is marked as fused. The system is evolved forward in time-steps of *δt* = 0.14s for a total time of 4 × 10^4^s, which is equivalent to 30*L*/*v*_*p*_. At each time step, the particles step in the appropriate direction a distance *vδt* (where *v* is the corresponding particle velocity) and undergo a transition event with probability 1 − *e*^−*kδt*^ (where *k* is the rate for that particular state transition).

## Acknowledgements

This research was supported by NIH grant R01 NS060698 to E.L.F.H., NSF Graduate Research Fellowship (DGE-1845298) to S.E.C., NSF CAREER grant (1848057) to EFK and a Cottrell Scholar Award to EFK. The authors declare no competing financial interests. We thank Andrea Stavoe, Alex Boecker, Dan Dou, Anamika Agrawal, and Keaton Holt for insights and discussions.

## Author contributions

Sydney E. Cason, Resources, Data curation, Formal analysis, Validation, Investigation, Visualization, Methodology, Project administration, Writing—original draft and review/editing; Saurabh S. Mogre, Conceptualization, Resources, Formal analysis, Investigation, Software, Validation, Visualization, Methodology, Project administration, Writing—original draft and review/editing; Erika L.F. Holzbaur, Supervision, Funding acquisition, Project administration, Writing—original draft and review/editing Elena F. Koslover, Conceptualization, Supervision, Software, Funding acquisition, Project administration, Writing—original draft and review/editing

---

**Figure 1–Figure supplement 1.**
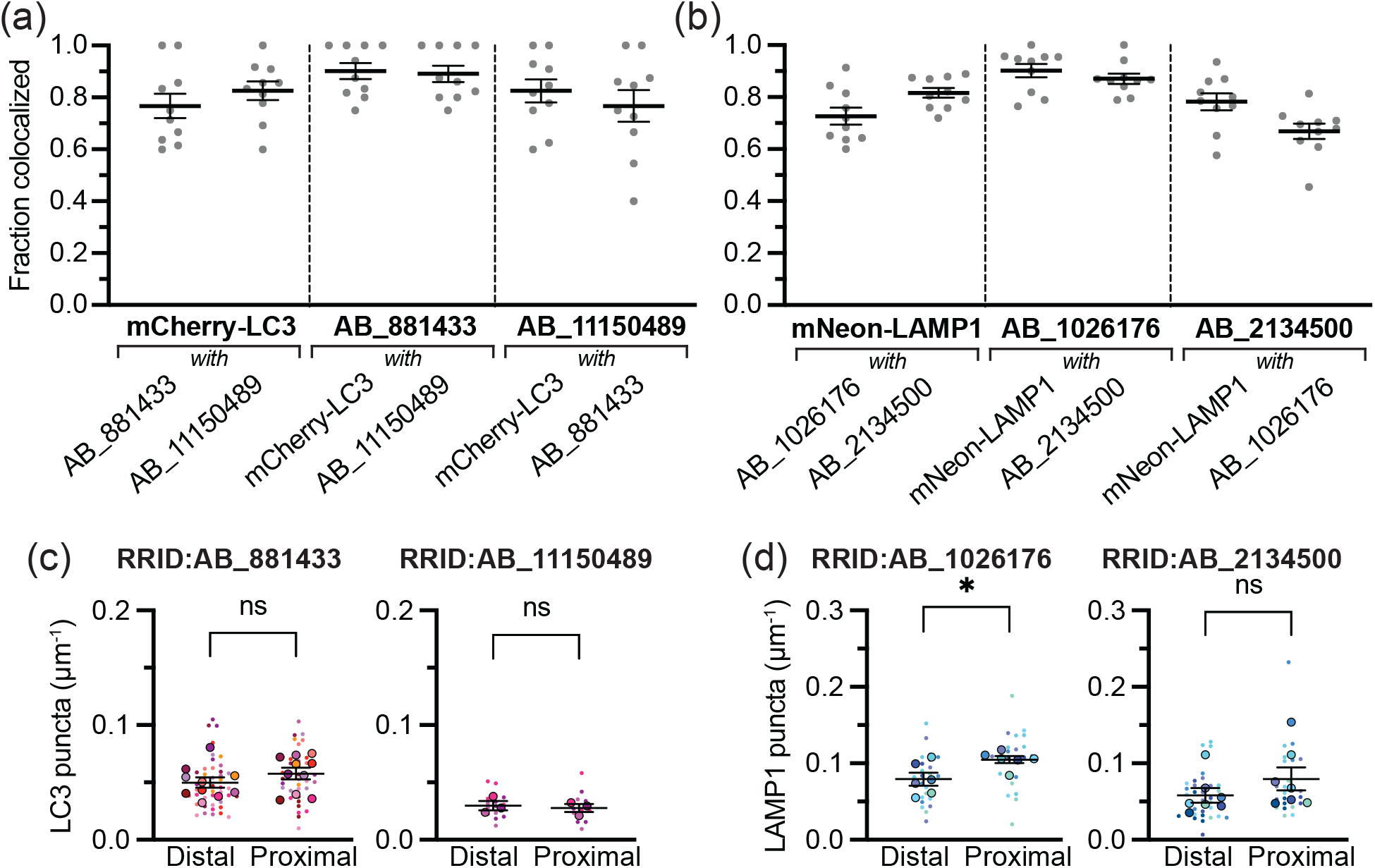
Antibody validation. **(a)** Colocalization between overexpressed mCherry-LC3 and both LC3 antibodies. *n* = 10 axons. **(b)** Colocalization between overexpressed mNeon-LAMP1 and both LAMP1 antibodies. *n* = 10 axons. **c** Linear density of LC3 puncta, probed with ab48394 (RRID:AB_881433; left; *n* = 38 − 43 axons; unpaired t test, *p* = 0.1199) or sc-376404 (RRID:AB_11150489; right; *n* = 12 − 14 axons; unpaired t test, *p* = 0.5997). **d** Linear density of LAMP1 puncta, probed with AF4800 (RRID:AB_1026176; left; *n* = 24 axons each; unpaired t test, *p* = 0.0174) or 1D4B (RRID:AB_2134500; right; *n* = 26 − 34 axons each; unpaired t test, *p* = 0.0497).

**Figure 1–Figure supplement 2.**
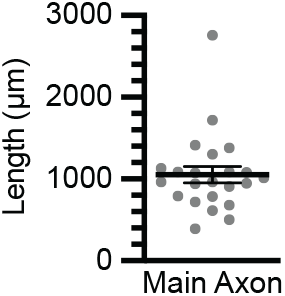
Axon length *in vitro*. Length of main axon (*n* = 23) as measured for primary hippocampal neurons at 7-10 days *in vitro*.

**Figure 1–Figure supplement 3.**
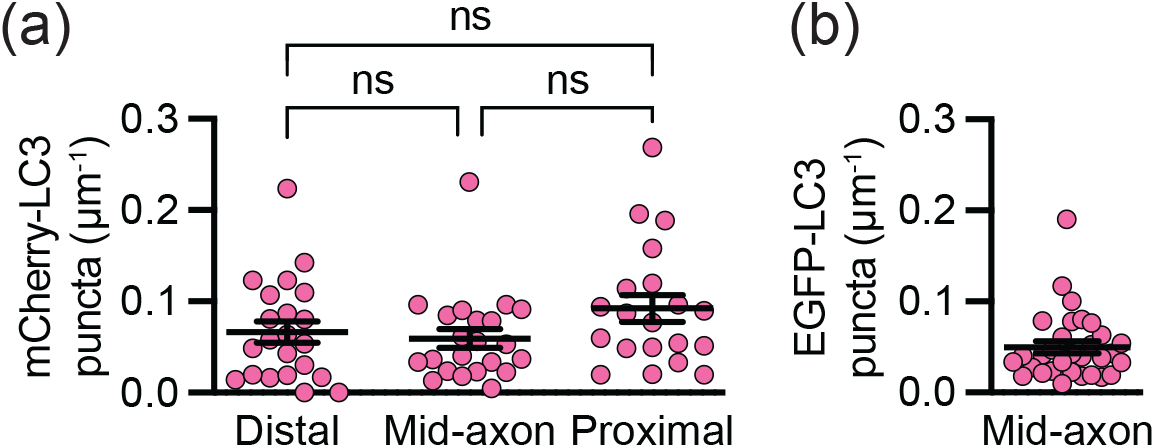
Linear LC3 density in live neuronal axons. **(a)** Linear density of overexpressed mCherry-LC3 in primary hippocampal neurons. *n* = 20-22 axons; one-way ANOVA, *p* = 0.1551. **(a)** Linear density of overexpressed EGFP-LC3 in iPSC-derived neurons. *n* = 31 axons.

**Figure 3–Figure supplement 1.**
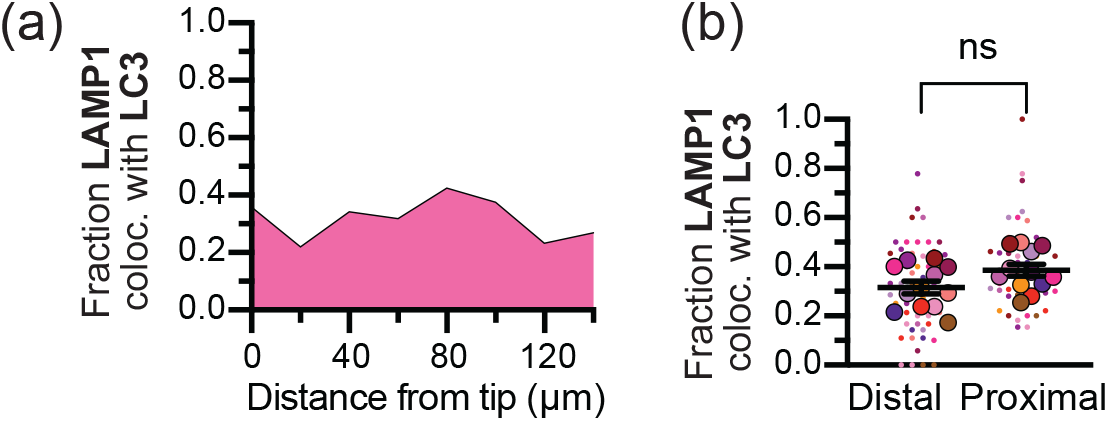
LAMP1 puncta colocalized with LC3. **(a)** Histogram showing the spatial distribution of LAMP1 puncta colocalized with LC3 in the distal axon. *n* = 773 puncta; 20 μm bins. **(b)** Comparison of the fraction LAMP1 puncta colocalized with LC3 in the distal and proximal axon. *n* = 12 trials; unpaired t test (*p* = 0.8411). Dashed line represents axon. ns, p > 0.05.

**Figure 7–Figure supplement 1.**
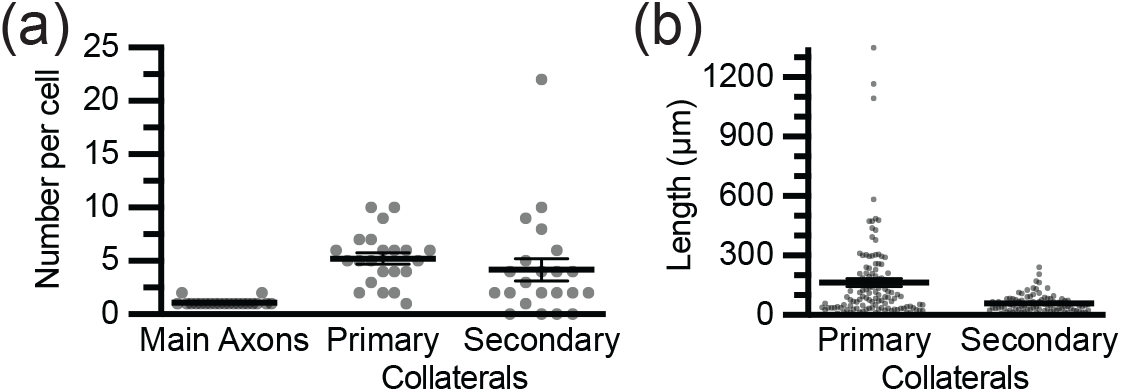
Number and length of axon collaterals. Primary collaterals are defined as branches off the main axon, while secondary collaterals are branches off the primary collaterals. Bifurcated main axons (whereby a cell may have 2+ axons) are distinguishable from primary collaterals based on branch angle (***Gallo, 2011***). **(a)** Number of axons or collaterals per cell. *n* = 22 cells. **(b)** Length of primary (*n* = 116) or secondary collaterals (*n* = 83).

**Figure 7–Figure supplement 2.**
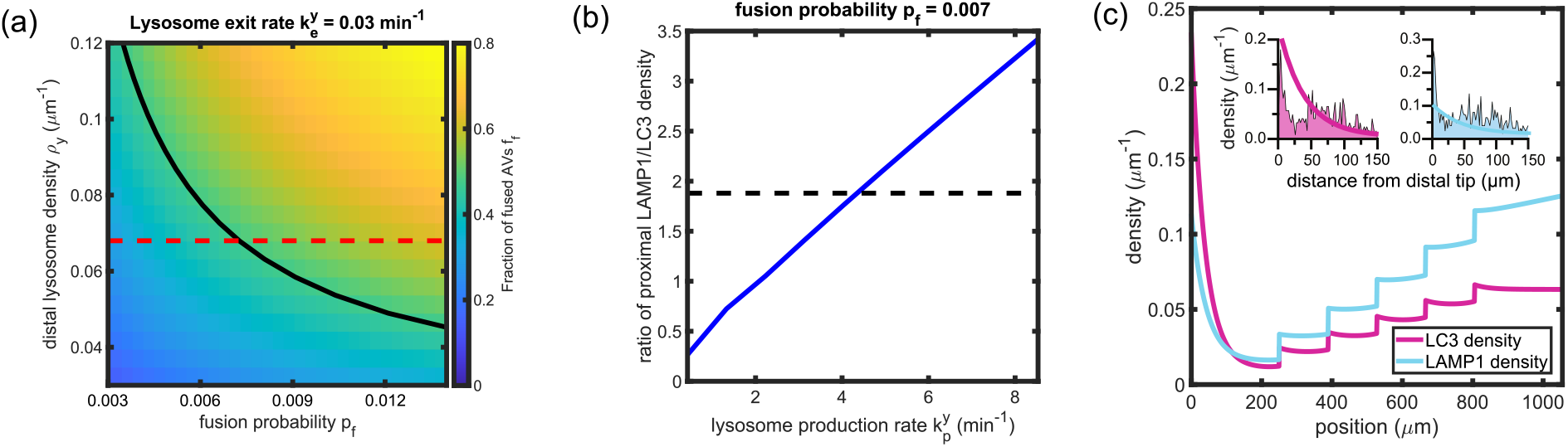
Modified model with unrestricted fusion predicts small number of fusion events per AV. **(a)** Fraction of AVs fused within the distal axon *f*_*f*_, plotted against the fusion probability *p*_*f*_, and the lysosome density in the distal region. The tip-exit rate for lysosomes 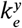 is set to 0.03 per minute. The solid black line denotes the measured value of *f*_*f*_ based on LC3+ puncta colocalized with LAMP1+ puncta in the distal axon. The dashed red line denotes the density of LAMP1+ puncta observed in the distal axon. **(b)** The ratio of the lysosome density to AV density in the proximal axon, plotted against the lysosome production rate. Dashed black line denotes the measured value determined by enumerating LAMP1+ and LC3+ puncta in the proximal axon. **(c)** The linear density of LC3+ puncta (magenta) and LAMP1+ puncta (cyan) along the axon. Insets show comparison to experimental data in the distal region, from Figure 1(d,f). **(d)** Spatial variation in the fraction of AVs fused at different positions along the axon (blue), and the average number of fusions per AV (red). The inset zooms into the distal region, overlaid with the observed distribution obtained by enumerating LC3+ puncta colocalized with LAMP1. All calculations were performed on the branched axon geometry shown in Figure 7a.

## References

Agrawal A, Koslover EF. Optimizing mitochondrial maintenance in extended neuronal projections. PLOS Computational Biology. 2021; 17(6):e1009073.

Agrotis A, Pengo N, Burden JJ, Ketteler R. Redundancy of human ATG4 protease isoforms in autophagy and LC3/GABARAP processing revealed in cells. Autophagy. 2019; 15(6):976–997.

Boecker CA, Goldsmith J, Dou D, Cajka GG, Holzbaur EL. Increased LRRK2 kinase activity alters neuronal autophagy by disrupting the axonal transport of autophagosomes. Current Biology. 2021;.

Boecker CA, Olenick MA, Gallagher ER, Ward ME, Holzbaur EL. ToolBox: Live Imaging of intracellular organelle transport in induced pluripotent stem cell-derived neurons. Traffic. 2020; 21(1):138–155.

Boyce WE, DiPrima RC, Meade DB. Elementary differential equations and boundary value problems. John Wiley & Sons; 2021.

Campbell TN, Choy FY. The effect of pH on green fluorescent protein: a brief review. Mol Biol Today. 2001; 2(1):1–4.

Cason SE, Carman PJ, Van Duyne C, Goldsmith J, Dominguez R, Holzbaur EL. Sequential dynein effectors regulate axonal autophagosome motility in a maturation-dependent pathway. Journal of Cell Biology. 2021; 220(7):e202010179.

Cheng XT, Xie YX, Zhou B, Huang N, Farfel-Becker T, Sheng ZH. Characterization of LAMP1-labeled nondegradative lysosomal and endocytic compartments in neurons. Journal of Cell Biology. 2018; 217(9):3127–3139.

Cheng XT, Zhou B, Lin MY, Cai Q, Sheng ZH. Axonal autophagosomes recruit dynein for retrograde transport through fusion with late endosomes. Journal of Cell Biology. 2015; 209(3):377–386.

Farfel-Becker T, Roney JC, Cheng XT, Li S, Cuddy SR, Sheng ZH. Neuronal soma-derived degradative lysosomes are continuously delivered to distal axons to maintain local degradation capacity. Cell reports. 2019; 28(1):51–64.

Ferguson SM. Axonal transport and maturation of lysosomes. Current opinion in neurobiology. 2018; 51:45–51.

Fu Mm, Nirschl JJ, Holzbaur EL. LC3 binding to the scaffolding protein JIP1 regulates processive dynein-driven transport of autophagosomes. Developmental cell. 2014; 29(5):577–590.

Fujita N, Hayashi-Nishino M, Fukumoto H, Omori H, Yamamoto A, Noda T, Yoshimori T. An Atg4B mutant hampers the lipidation of LC3 paralogues and causes defects in autophagosome closure. Molecular biology of the cell. 2008; 19(11):4651–4659.

Gallo G. The cytoskeletal and signaling mechanisms of axon collateral branching. Developmental neurobiology. 2011; 71(3):201–220.

Goldsmith J, Ordureau A, Harper JW, Holzbaur EL. Brain-derived autophagosome profiling reveals the engulfment of nucleoid-enriched mitochondrial fragments by basal autophagy in neurons. Neuron. 2022;.

Gowrishankar S, Yuan P, Wu Y, Schrag M, Paradise S, Grutzendler J, De Camilli P, Ferguson SM. Massive accumulation of luminal protease-deficient axonal lysosomes at Alzheimer’s disease amyloid plaques. Proceedings of the National Academy of Sciences. 2015; 112(28):E3699–E3708.

Hara T, Nakamura K, Matsui M, Yamamoto A, Nakahara Y, Suzuki-Migishima R, Yokoyama M, Mishima K, Saito I, Okano H, et al. Suppression of basal autophagy in neural cells causes neurodegenerative disease in mice. Nature. 2006; 441(7095):885–889.

Hill SE, Kauffman KJ, Krout M, Richmond JE, Melia TJ, Colón-Ramos DA. Maturation and clearance of autophagosomes in neurons depends on a specific cysteine protease isoform, ATG-4.2. Developmental cell. 2019; 49(2):251–266.

Itakura E, Kishi-Itakura C, Mizushima N. The hairpin-type tail-anchored SNARE syntaxin 17 targets to autophagosomes for fusion with endosomes/lysosomes. Cell. 2012; 151(6):1256–1269.

Johnson DE, Ostrowski P, Jaumouillé V, Grinstein S. The position of lysosomes within the cell determines their luminal pH. Journal of Cell Biology. 2016; 212(6):677–692.

Kalil K, Dent EW. Branch management: mechanisms of axon branching in the developing vertebrate CNS. Nature Reviews Neuroscience. 2014; 15(1):7–18.

Kauffman KJ, Yu S, Jin J, Mugo B, Nguyen N, O’Brien A, Nag S, Lystad AH, Melia TJ. Delipidation of mammalian Atg8-family proteins by each of the four ATG4 proteases. Autophagy. 2018; 14(6):992–1010.

Koltun B, Ironi S, Gershoni-Emek N, Barrera I, Hleihil M, Nanguneri S, Sasmal R, Agasti SS, Nair D, Rosenblum K. Measuring mRNA translation in neuronal processes and somata by tRNA-FRET. Nucleic Acids Research. 2020 01; 48(6):e32–e32. https://doi.org/10.1093/nar/gkaa042, doi: 10.1093/nar/gkaa042.

Komatsu M, Waguri S, Chiba T, Murata S, Iwata Ji, Tanida I, Ueno T, Koike M, Uchiyama Y, Kominami E, et al. Loss of autophagy in the central nervous system causes neurodegeneration in mice. Nature. 2006; 441(7095):880–884.

Lee S, Sato Y, Nixon RA. Lysosomal Proteolysis Inhibition Selectively Disrupts Axonal Transport of Degradative Organelles and Causes an Alzheimer’s-Like Axonal Dystrophy. Journal of Neuroscience. 2011; 31(21):7817–7830. https://www.jneurosci.org/content/31/21/7817, doi: 10.1523/JNEUROSCI.6412-10.2011.

Li Y, Cheng X, Li M, Wang Y, Fu T, Zhou Z, Wang Y, Gong X, Xu X, Liu J, et al. Decoding three distinct states of the Syntaxin17 SNARE motif in mediating autophagosome–lysosome fusion. Proceedings of the National Academy of Sciences. 2020; 117(35):21391–21402.

Li Y, Wang X, Li M, Yang C, Wang X. M05B5. 4 (Lysosomal phospholipase A2) promotes disintegration of autophagic vesicles to maintain C. elegans development. Autophagy. 2021; (just-accepted).

Lie PP, Yang DS, Stavrides P, Goulbourne CN, Zheng P, Mohan PS, Cataldo AM, Nixon RA. Post-Golgi carriers, not lysosomes, confer lysosomal properties to pre-degradative organelles in normal and dystrophic axons. Cell reports. 2021; 35(4):109034.

Ma X, Godar RJ, Liu H, Diwan A. Enhancing lysosome biogenesis attenuates BNIP3-induced cardiomyocyte death. Autophagy. 2012; 8(3):297–309.

Maday S, Holzbaur EL. Autophagosome biogenesis in primary neurons follows an ordered and spatially regulated pathway. Developmental cell. 2014; 30(1):71–85.

Maday S, Wallace KE, Holzbaur EL. Autophagosomes initiate distally and mature during transport toward the cell soma in primary neurons. Journal of Cell Biology. 2012; 196(4):407–417.

Maehr R, Hang HC, Mintern JD, Kim YM, Cuvillier A, Nishimura M, Yamada K, Shirahama-Noda K, Hara-Nishimura I, Ploegh HL. Asparagine endopeptidase is not essential for class II MHC antigen presentation but is required for processing of cathepsin L in mice. The Journal of Immunology. 2005; 174(11):7066–7074.

Martens S, Fracchiolla D. Activation and targeting of ATG8 protein lipidation. Cell discovery. 2020; 6(1):1–11.

MATLAB. version 9.10.0 (R2021a). Natick, Massachusetts: The MathWorks Inc.; 2021.

Maxson ME, Abbas YM, Wu JZ, Plumb JD, Grinstein S, Rubinstein JL. Detection and quantification of the vacuolar H+ ATPase using the Legionella effector protein SidK. Journal of Cell Biology. 2022; 221(3):e202107174.

Misgeld T, Schwarz TL. Mitostasis in neurons: maintaining mitochondria in an extended cellular architecture. Neuron. 2017; 96(3):651–666.

Mogre SS, Christensen JR, Niman CS, Reck-Peterson SL, Koslover EF. Hitching a ride: mechanics of transport initiation through linker-mediated hitchhiking. Biophysical journal. 2020; 118(6):1357–1369.

Mogre SS, Christensen JR, Reck-Peterson SL, Koslover EF. Optimizing microtubule arrangements for rapid cargo capture. Biophysical Journal. 2021; 120(22):4918–4931.

Pankiv S, Clausen TH, Lamark T, Brech A, Bruun JA, Outzen H, Øvervatn A, Bjørkøy G, Johansen T. p62/SQSTM1 binds directly to Atg8/LC3 to facilitate degradation of ubiquitinated protein aggregates by autophagy. Journal of biological chemistry. 2007; 282(33):24131–24145.

Pantazis CB, Yang A, Lara E, McDonough JA, Blauwendraat C, Peng L, Oguro H, Zou J, Sebesta D, Pratt G, et al. A reference induced pluripotent stem cell line for large-scale collaborative studies. bioRxiv. 2021;.

Roney JC, Li S, Farfel-Becker T, Huang N, Sun T, Xie Y, Cheng XT, Lin MY, Platt FM, Sheng ZH. Lipid-mediated motor-adaptor sequestration impairs axonal lysosome delivery leading to autophagic stress and dystrophy in Niemann-Pick type C. Developmental Cell. 2021; 56(10):1452–1468.

Saleeb RS, Kavanagh DM, Dun AR, Dalgarno PA, Duncan RR. A VPS33A-binding motif on syntaxin 17 controls autophagy completion in mammalian cells. Journal of Biological Chemistry. 2019; 294(11):4188–4201.

Shen Q, Shi Y, Liu J, Su H, Huang J, Zhang Y, Peng C, Zhou T, Sun Q, Wan W, et al. Acetylation of STX17 (syntaxin 17) controls autophagosome maturation. Autophagy. 2021; 17(5):1157–1169.

Shibutani ST, Yoshimori T. A current perspective of autophagosome biogenesis. Cell research. 2014; 24(1):58–68.

Stavoe AK, Gopal PP, Gubas A, Tooze SA, Holzbaur EL. Expression of WIPI2B counteracts age-related decline in autophagosome biogenesis in neurons. Elife. 2019; 8:e44219.

Stavoe AK, Hill SE, Hall DH, Colón-Ramos DA. KIF1A/UNC-104 transports ATG-9 to regulate neurodevelopment and autophagy at synapses. Developmental cell. 2016; 38(2):171–185.

Stavoe AK, Holzbaur EL. Autophagy in neurons. Annual review of cell and developmental biology. 2019; 35:477–500.

Tsuboyama K, Koyama-Honda I, Sakamaki Y, Koike M, Morishita H, Mizushima N. The ATG conjugation systems are important for degradation of the inner autophagosomal membrane. Science. 2016; 354(6315):1036–1041.

Williams AH, O’donnell C, Sejnowski TJ, O’leary T. Dendritic trafficking faces physiologically critical speed-precision tradeoffs. elife. 2016; 5:e20556.

Wong YC, Holzbaur EL. Autophagosome dynamics in neurodegeneration at a glance. Journal of cell science. 2015; 128(7):1259–1267.

Yin Z, Pascual C, Klionsky DJ. Autophagy: machinery and regulation. Microbial cell. 2016; 3(12):588.

Yu L, McPhee CK, Zheng L, Mardones GA, Rong Y, Peng J, Mi N, Zhao Y, Liu Z, Wan F, et al. Termination of autophagy and reformation of lysosomes regulated by mTOR. Nature. 2010; 465(7300):942–946.

